# Iron-deficiency and estrogen are associated with ischemic stroke by up-regulating transferrin to induce hypercoagulability

**DOI:** 10.1101/646109

**Authors:** Xiaopeng Tang, Mingqian Fang, Ruomei Cheng, Zhiye Zhang, Yuming Wang, Chuanbin Shen, Yajun Han, Qiumin Lu, Yingrong Du, Yingying Liu, Zhaohui Sun, Liping Zhu, James Mwangi, Min Xue, Chengbo Long, Ren Lai

**Author notes:** These authors contributed equally to this work (Xiaopeng Tang, Mingqian Fang, Ruomei Cheng, Zhiye Zhang, Yuming Wang, Chuanbin Shen). **Corresponding authors** Address correspondence to: Kunming Institute of Zoology, Chinese Academy of Sciences, Kunming 650223, Yunnan, China; (RL).

## Abstract

In the accompanying manuscript, transferrin has been demonstrated to maintain coagulation balance by interacting with clotting factors, suggesting that elevated transferrin causes thromboembolic diseases and factors up-regulating transferrin is associated with thrombosis. Here we show that transferrin and transferrin-thrombin/FXIIa complexes are elevated in plasma and cerebrospinal fluid of ischemic stroke (IS) patients with iron-deficiency anemia (IDA) history, IDA patients and venous thromboembolism patients using combined oral contraceptives (CC) as well as ID mice, suggesting an association of transferrin up-regulation with ID and CC. ID and estrogen up-regulated transferrin through hypoxia and estrogen response elements located at transferrin gene enhancer and promoter region, respectively. ID, exogenous transferrin/estrogen administration or transferrin over-expression promoted hypercoagulability and aggravated IS, while anti-transferrin antibody, transferrin knockdown or designed peptide inhibitors interfering transferrin-thrombin/FXIIa interaction exerted anti-IS effects in *vivo*. Collectively, the results reveal that factors (i.e., ID and CC) up-regulating transferrin are risk factors of thromboembolic diseases.

## Introduction

Ischemic stroke (IS) is typically caused by blockage of a blood vessel (Hacke et al., 2008), counting for approximately 87% of strokes (Kulcsar et al., 2014; Talwalkar and Uddin, 2015). Briefly, IS occurs upon the severe reduction and interruption of blood flow (ischemia) within the cerebral circulation induced by vascular occlusion(Maeda et al., 1999; Novakovic et al., 2009; Weimar et al., 2009). IS is the second-leading cause of death and the most important cause of permanent disability worldwide (Feigin et al., 2014; Feigin et al., 2016). Two acquired conditions or environment factors including iron deficiency (ID) and combined oral contraceptives (CC) are known to be associated with an increased risk of thromboembolism such as IS (Akins et al., 1996; Batur Caglayan et al., 2013; Belman et al., 1990; Benedict et al., 2004; Coutinho et al., 2015; Devuyst et al., 2000; Hartfield et al., 1997; Kinoshita et al., 2006; Maguire et al., 2007; Makhija et al., 2008; Nagai et al., 2005; Roach et al., 2015) and venous thromboembolism (VTE) (Coutinho et al., 2015; Jick et al., 2000; Kemmeren et al., 2001; Lewis and Heinemann, 1997; Stegeman et al., 2013; Vandenbroucke et al., 2001).

ID is the most common nutritional disorder affecting more than two billion people worldwide(McLean et al., 2009), and iron-deficiency anemia (IDA) remains the foremost cause of anemia (Kassebaum et al., 2014). In 1983, Alexander reported that IDA is associated with hemiparesis and aphasia(Alexander, 1983), which firstly drew attention to the correlation between IDA and thrombotic complications. Over the last few years, association of ID/IDA and thrombophilia has been increasingly recognized (Akins et al., 1996; Batur Caglayan et al., 2013; Belman et al., 1990; Benedict et al., 2004; Coutinho et al., 2015; Devuyst et al., 2000; Hartfield et al., 1997; Kinoshita et al., 2006; Maguire et al., 2007; Makhija et al., 2008; Nagai et al., 2005). Various kinds of thrombotic diseases including central retinal vein occlusion (Nagai et al., 2005), cerebral venous sinus thrombosis (Belman et al., 1990; Kinoshita et al., 2006), cerebral sinovenous thrombosis (Benedict et al., 2004), and carotid artery thrombus(Akins et al., 1996) were observed to be associated with IDA. In addition, numbers of cases of embolic stroke (Hartfield et al., 1997; Maguire et al., 2007; Makhija et al., 2008; Roshal, 2016) and IS (Chang et al., 2013; Ganesan et al., 2003; Munot et al., 2011) have been reported to be associated with ID/IDA. Population-based studies found that IDA is more frequent in cerebral venous thrombosis cases (27.0%) than in healthy controls (6.5%)(Coutinho et al., 2015). Meta-analysis shows that the risk of myocardial infarction or ischemic stroke is 1.6-fold increase in women using CC (Roach et al., 2015). CC are also associated with an increased risk of venous thromboembolism (VTE) (Jick et al., 2000; Kemmeren et al., 2001; Lewis and Heinemann, 1997; Stegeman et al., 2013; Vandenbroucke et al., 2001). In 2017, Practice Committee of the American Society for Reproductive Medicine also issues that CC increase the risk of VTE (Med, 2017). All of these previous data suggest that ID and CC are associated with thrombotic diseases and IS, but the underlying mechanisms are not known. Given that transferrin (Tf) interacts with and potentiate FXIIa/thrombin and blocks antithrombin III (AT-III)’s inactivation on coagulation proteases to induce hypercoagulability as reported in the accompanying manuscript, we here investigate if Tf mediates the association between ID, CC and thrombosis.

## Results

### Elevated Tf, Tf-thrombin/FXIIa complexes in plasma and cerebrospinal fluid (CSF) of IS, IDA and VTE patients

As illustrated in Figure 1A, elevated Tf was detected in the plasma of IS patients with IDA history, IDA patients and VTE patients using CC by enzyme linked immunosorbent assay (ELISA). The average Tf concentration in plasma of IS patients (n = 453, male 262; female 191), IDA patients (n = 454, male 228; female 226) and VTE patients using CC (n = 30, female) was 4.497 ± 1.57 mg/ml, 4.146 ± 2.06 mg/ml and 5.27 ± 0.94 mg/ml, respectively, while that in healthy individuals (n = 341, male 140; female 201) was 2.820 ± 0. 55 mg/ml. Western blot also exhibited an elevated Tf level in plasma of those patients (Figures 1B and 1C). In addition, we found that Tf in cerebrospinal fluid of IS patients with IDA history (n = 40, male 22; female 18) was 20.11 ± 5.41 mg/L, which was ∼0.5-fold greater than that (13.7 ± 2.70 mg/L) in healthy controls (n = 40, male 20; female 20) (Figures 1D-1F).

**Figure 1.**
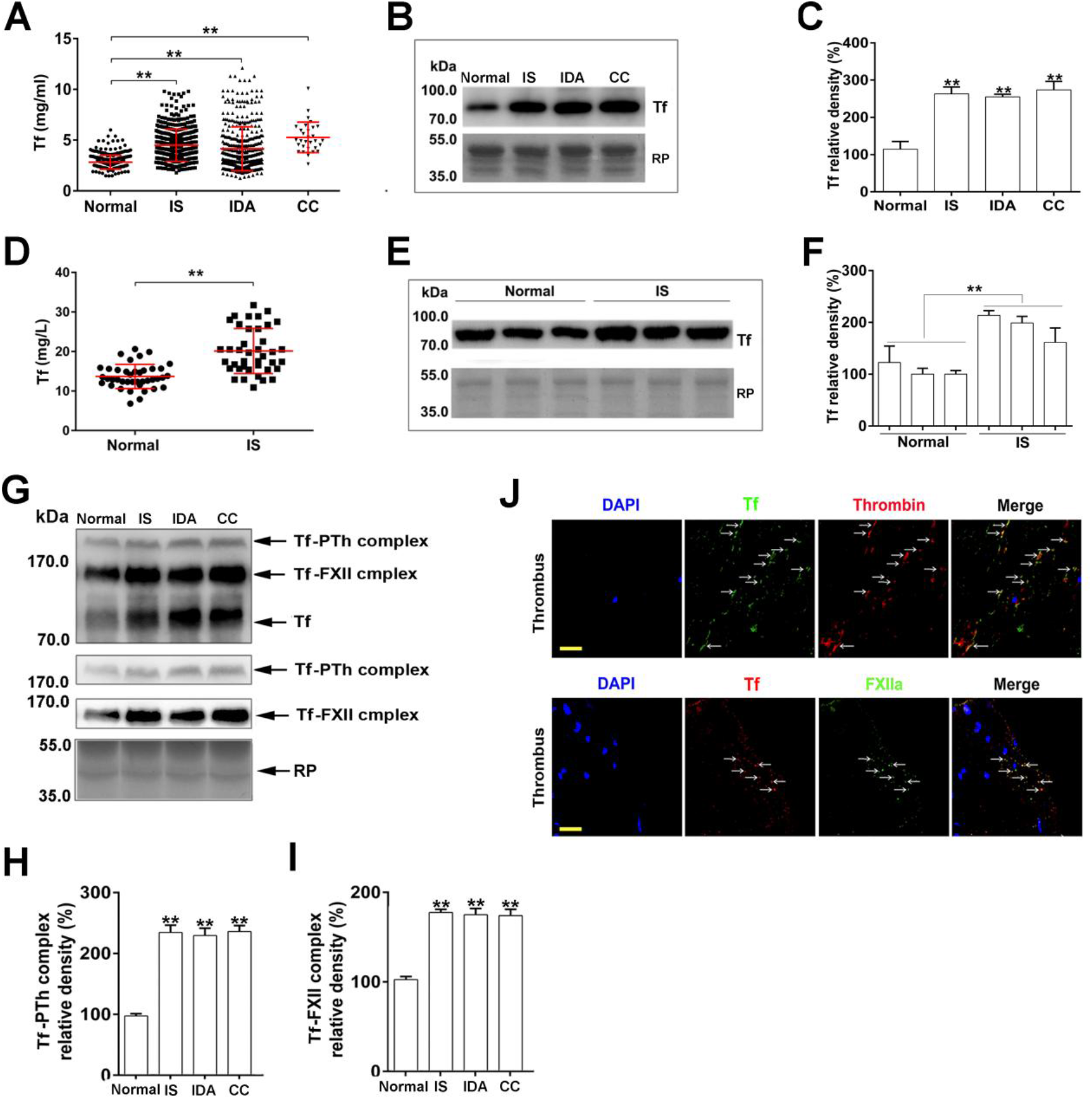
Elevated transferrin and transferrin-thrombin/FXIIa complexes are found in plasma or cerebrospinal fluid of ischemic stroke, iron-deficiency anemia and venous thromboembolism patients. **(A)** The concentration of Tf in plasma of IS patients, IDA patients, VTE patients using CC (CC) and volunteers (normal) were determined by ELISA. Western blot **(B)** and quantification **(C)** of Tf in plasma samples from IS patients, IDA patients, VTE patients using CC and volunteers. **(D)** The level of Tf in CSF of IS patients and controls (normal). Western blot **(E)** and quantification **(F)** of Tf in CSF samples of IS patients and controls. **(G)** After pre-treated by BS^3^, the plasma of IS patients, IDA patients, VTE patients using CC and normal controls were utilized to detect the formation of Tf-prothrombin and Tf-FXII complexes by western blot analysis using anti-Tf antibody. The formed complexes were indicated by arrows. Tf-prothrombin and Tf-FXII complexes in plasma were further confirmed by using anti-thrombin and anti-FXII antibody after stripping the PVDF membranes, respectively. Quantifications of Tf-prothrombin (PTh) **(H)** and Tf-FXII complexes **(I). (J)** Clots from acute IS patients were labeled with either anti-Tf (green) or anti-thrombin antibody (red) to detect the presence of Tf-thrombin complex (top), or labeled with either anti-Tf (red) or anti-FXIIa antibody (green) to detect the presence of Tf-FXIIa complex (bottom). Cell nuclei were labeled by DAPI. Arrows indicate Tf-thrombin- or Tf-FXIIa-positive structures. Scale bar represents 30 μm. Images were representative of at least three independent experiments. **(B, E** and **G)** Red Ponceau (RP)-stained blot (below) is the loading control. Data represent mean ± SD, \*\**p* < 0.01 by **(A** and **D)** unpaired t-test. Data represent mean ± SD of six independent experiments of 18 patients, respectively, \*\**p* < 0.01 by **(B** and **E)** unpaired t-test. Data represent mean ± SD of five independent experiments of 10 patients, respectively, \*\**p* < 0.01 by **(G)** unpaired t-test.

Western blot results demonstrated 3 bands recognized by anti-Tf antibody in the Bis (sulfosuccinimidyl) suberate (BS^3^) crosslinked plasma of IS patients with IDA history, IDA patients and VTE patients using CC (Figure 1G, the top panel). One of them was Tf, and the other 2 bands were Tf-prothrombin/FXII complexs, which are confirmed by antibody against prothrombin and FXII (Figure 1G, the 2 medium panels), respectively. Notably, the amounts of both the Tf-thrombin and Tf-FXIIa complexes in IS patients, IDA patients and VTE patients using CC were significantly higher than those in normal controls (Figures 1H-1I). To determine whether the Tf-thrombin or Tf-FXIIa complexes were present in thrombus, the colocalization of Tf with thrombin or FXIIa in thrombus was detected by confocal microscopy. As shown in Figure 1J (top), most Tf-positive deposits (green) are associated with thrombin (red), indicating the formation of Tf-thrombin complex *in vivo*. Colocalization of Tf (red) and FXIIa (green) by immunofluorescence (Figure 1J, bottom) also confirmed the formation of Tf-FXIIa complex in the thrombus.

### ID up-regulates Tf expression *in vitro*

We next investigated if ID affects Tf expression by using a mouse normal liver cell line BNL CL.2. *Tf* mRNA expression (Figure 2A) was up-regulated by iron chelator desferrioxamine (DFO) in a dose-dependent manner. Tf protein was also up-regulated by DFO in a dose-dependent manner by ELISA (Figure 2B) and western blot (Figures 2C and 2D) analysis, respectively. It has been previously reported that hypoxia-inducible factor-1 (HIF-1) confers low oxygen up-regulation of Tf gene expression by binding to the two binding sites of HIF-1 present in the Tf enhancer (Rolfs et al., 1997). As illustrated in Figures 2E and 2F, DFO administration up-regulated HIF-1α in a dose-dependent manner, while DFO showed no significant effect on Tf expression when the binding sites of HIF-1 present in the enhancer region of Tf gene were mutated (Figure 2G).

**Figure 2.**
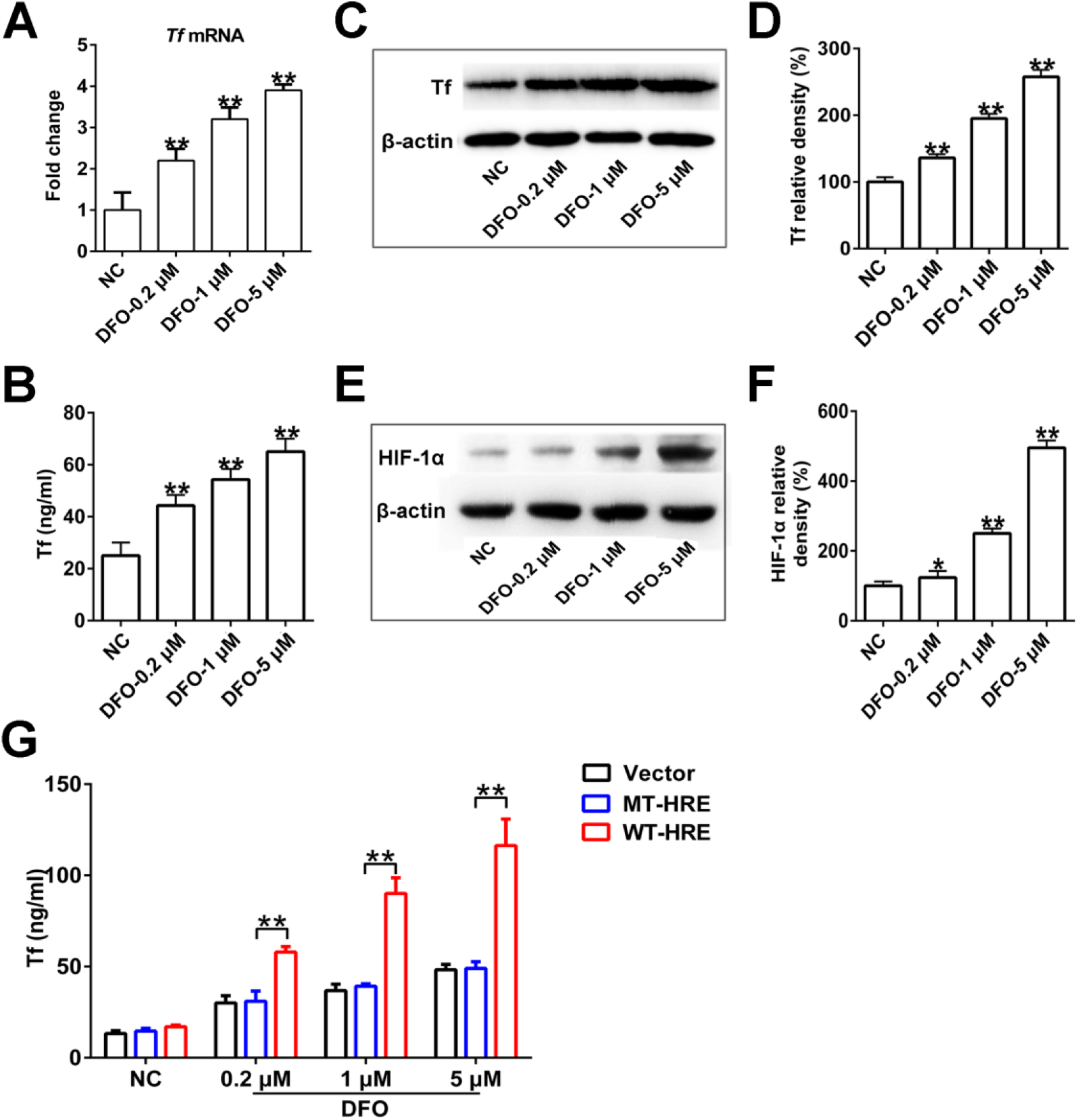
Transferrin is up-regulated by iron-deficiency *in vitro*. After treatment by iron chelator DFO, *Tf* RNA expression in BNL CL.2 cells were analyzed by quantitative real-time polymerase chain reaction (qRT-PCR) **(A)**. Tf protein in BNL CL.2 cells was analyzed by ELISA **(B)** and western blot **(C)**, respectively. **(D)** Quantitative analysis of western blot in **(C)**. BNL CL.2 cells were treated by different concentrations of DFO, HIF-1α was analyzed by western blot **(E)** and the quantification of “E” was shown **(F)**. **(G)** Tf level in supernatant of HepG2 cells transfected by Tf expression plasmid of wild-type HRE (WT-HRE) or mutated HRE (MT-HRE) was analyzed by ELISA after treatment by DFO. Data represent mean ± SD of six independent experiments, \*\**p* < 0.01 by one-way ANOVA with Dunnett’s post hoc test.

### ID induces Tf up-regulation to promote hypercoagulability *in vivo*

To elucidate the relationship of Tf and ID, mice were subjected to a low-iron diet or high-iron diet. After 4 weeks, a substantial increase in Tf plasma level was observed in low-iron diet induced mice compared with normal controls or high-iron diet induced mice (Figure 3A). The elevated Tf was associated with typical characteristics of microcytic hypochromic anemia, such as decreased mean corpusular volume (MCV) and mean corpusular hemoglobin (MCH), etc (Table S4). Furthermore, a significant reduction in activated partial thromboplastin time (APTT) (Figure 3B), prothrombin time (PT) (Figure 3C), and plasma recalcification time (Figures 3D and 3E) was observed in low-iron diet induced mice by comparison with the control mice or high-iron diet induced mice. Moreover, an obvious decrease in tail-bleeding time was observed in low-iron diet induced mice (Figure 3F). Importantly, anti-Tf antibody-treatment reversed the reduction of APTT, PT, plasma recalcification time and tail-bleeding time (Figures 3B-3F), while control IgG did not reverse the reduction of APTT, PT, plasma recalcification time and tail-bleeding time (Figures 3B-3F). As illustrated in Figure 3G, decreased blood flow was observed in the carotid artery of low-iron diet induced mice by comparison with the control mice or high-iron diet induced mice. Anti-Tf antibody-treatment reversed the reduction of blood flow (Figure 3G). The results demonstrate that Tf is up-regulated upon ID/IDA and induces hypercoagulability and thrombus formation.

**Figure 3.**
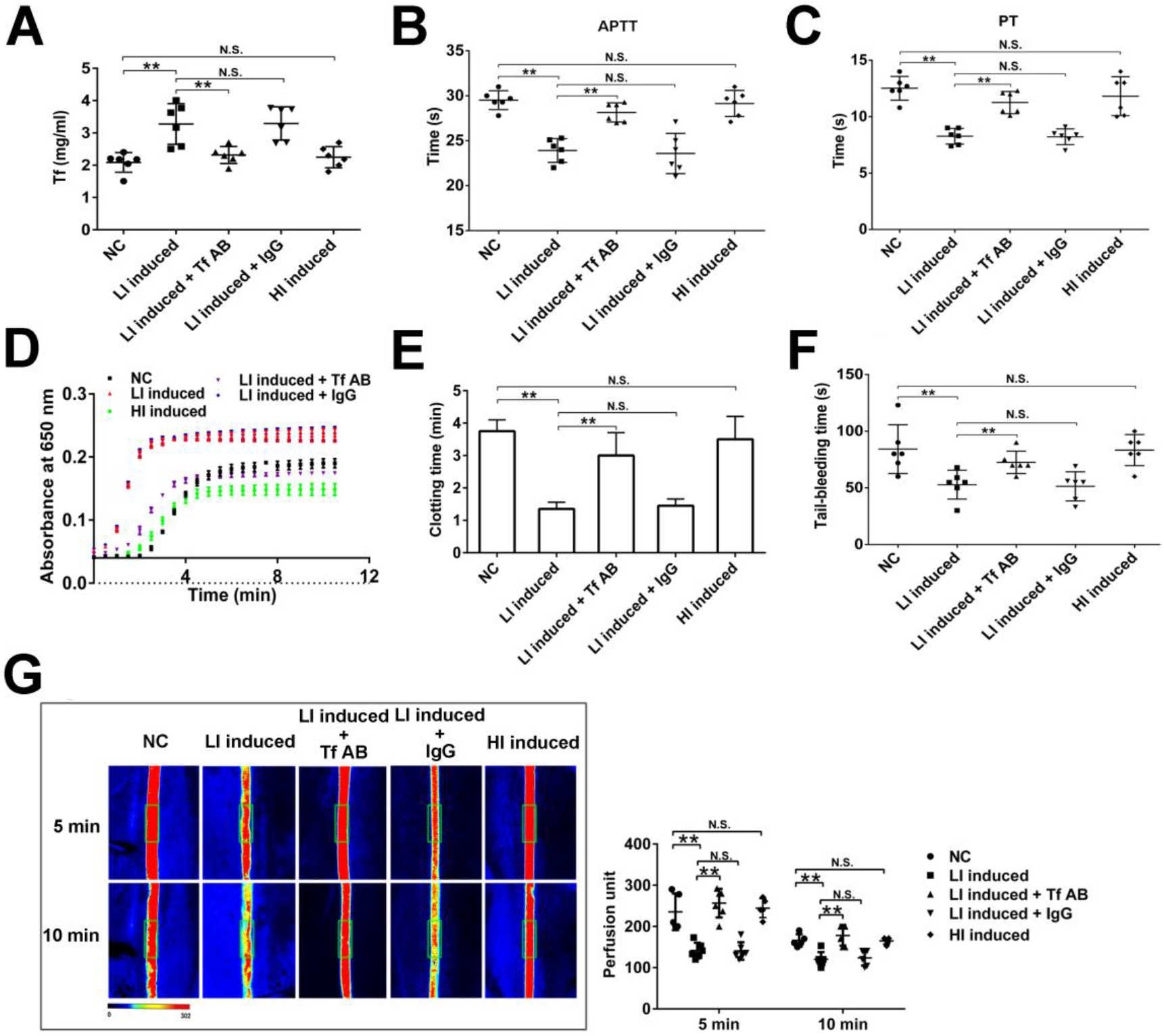
Iron-deficiency induces hypercoagulability and thrombus formation by up-regulating transferrin in mice. **(A)** Plasma Tf concentration in mice fed by normal diet (NC), low-iron diet (LI induced) or high-iron diet (HI induced) was determined by ELISA, respectively. APTT **(B)**, PT **(C)**, plasma recalcification time **(D)** and **(E)** (the clotting time was calculated by measuring the time to half maximal increase in absorbance), tail-bleeding time **(F)** were determined by using the mice fed a normal diet (NC), low-iron diet (LI induced) or high-iron diet (HI induced) alone, or from the mice treated with anti-Tf antibody (LI induced + Tf AB) or control IgG (LI induced + IgG) when fed a low-iron diet. **(G)** Representative images of carotid artery blood flow (left) in FeCl_3_-treated all groups of mice by laser speckle perfusion imaging, and the region of interest (the rectangle in green) was placed in the carotid artery to quantify blood flow change. Relative blood flow in the region of interest was shown (right). Animal experiments were repeated three independent times. Data represent mean ± SD (n = 5-6), \*\**p* < 0.01 by one-way ANOVA with Dunnett’s post hoc test. N.S.: no significance.

### Estrogen up-regulates Tf expression *in vitro*

We next investigated estrogen’s effect on Tf expression by using a mouse normal liver cell line BNL CL.2. Tf protein was up-regulated by estrogen in a dose-dependent manner by both ELISA (Figure 4A) and western blot (Figures 4B and 4C) analysis. Previous study has indicated that the estrogen response elements (EREs) located at human Tf gene promoter region is associated with a GGACA(N)_3_TGGCC motif, which binds estrogen receptor α (Vyhlidal et al., 2002). We found a similar motif (GGAGAGGATGGCC) at 5’ promoter of mouse Tf gene between −1021 and −1009. As illustrated in Figure 4D, Tf up-regulation induced by estrogen in BNL CL.2 cells was significantly inhibited when the corresponding ERE in promoter region of Tf gene was mutated.

**Figure 4.**
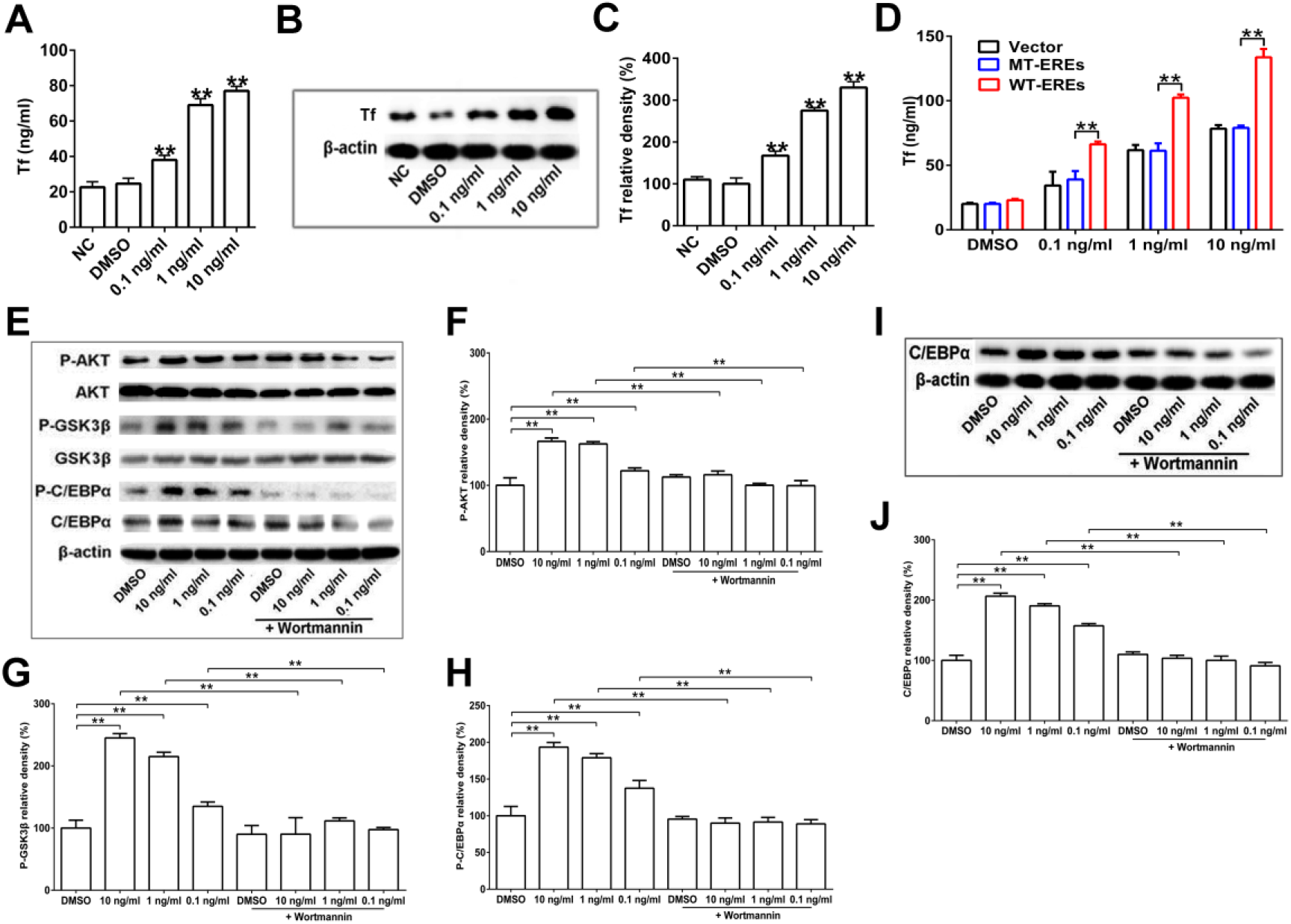
Transferrin is up-regulated by estrogen *in vitro*. After treatment by estrogen, Tf expression in BNL CL.2 cells were analyzed by ELISA **(A)** and western blot (**B** and **C**), respectively. **(D)** Tf level in supernatant of BNL CL.2 cells transfected by Tf expression plasmid of wild-type EREs (WT-EREs) or mutated EREs (MT-EREs) was analyzed by ELISA. **(E)** BNL CL.2 cells were treated by estrogen in the presence or absence of wortmanmin for 6 h, total and phosphorylated AKT, GSK3β, and C/EBPα were analyzed by western blot, respectively. β-actin was used as the loading control. The relative density of P-AKT **(F)**, P-GSK3β **(G)** and P-C/EBPα **(H)** was quantified. Western blot **(I)** and quantification **(J)** of C/EBPα in BNL CL.2 cells treated by estrogen alone or in the presence of wortmanmin for 24 h. β-actin was the loading control. P-AKT: phospho-Akt; P-GSK3β: phospho-GSK3β; P-C/EBPα: phospho-C/EBPα. Data represent mean ± SD of five independent experiments, \*\**p* < 0.01 by one-way ANOVA with Dunnett’s post hoc test.

CCAAT/enhancer binding protein-α (C/EBPα) is a key transcription factor involved in numerous genes’ expression by interacting with the regulatory sequences presented in the promoter and enhancer regions of target genes (Ramji and Foka, 2002). AKT-dependent phosphorylation of glycogen synthase kinase-3β (GSK3β) has been reported to up-regulate the activity of C/EBPα (Chen et al., 2016). As illustrated in Figures 4E and 4F, AKT was activated by estrogen in a dose-dependent manner. Activated AKT in turn phosphorylated and inactivated GSK3β (Figure 4G). GSK3β phosphorylation was increased by ∼30, 115 and 146% after 2-h treatment with 0.1, 1 and 10 ng/ml of estrogen, respectively (Figure 4G). About 45, 80 and 95% elevation of C/EBPα phosphorylation was also observed after treatment with 0.1, 1 and 10 ng/ml of estrogen, respectively (Figures 4E and 4H). Furthermore, C/EBPα expression was elevated after 24-h treatment with estrogen (Figures 4I and 4J). The phosphorylation of AKT, GSK3β and C/EBPα, and expression of C/EBPα induced by estrogen were significantly abrogated by phosphatidylinositol 3-kinase (PI3K) inhibitor wortmannin (10 nM) (Figures 4E-4J), suggesting that PI3K/AKT signaling pathway is involved in Tf expression induced by estrogen. Together, all these data indicate that estrogen up-regulates Tf expression through the ERE and AKT activation.

### Estrogen induces Tf up-regulation to promote hypercoagulability *in vivo*

As illustrated in Figure 5B, Tf in plasma of both male and female mice was up-regulated by oral gavage administration of estrogen, which was in accordance with estrogen elevation in mouse plasma (Figure 5A). Compared with 2.4 mg/ml in the plasma of male control mice, Tf concentration was elevated to 3.0, 3.6, and 4.2 mg/ml by 0.2, 1 and 5 mg/kg estrogen administration, respectively (Figure 5B). The similar situation was also observed in female mice (Figure 5B). We then determined whether the association of estrogen and hypercoagulability is dependent on Tf. After gavage administration with estrogen (1 mg/kg) for one week (long-term administration), the concentration of Tf in mice plasma was ∼3.9 mg/ml, while that in controls was ∼2.3 mg/ml (Figure 5C). After one-week estrogen administration, APTT and PT were determined to elucidate whether estrogen treatment induced hypercoagulability. As illustrated in Figure 5D, APTT in control mice was ∼29 sec, while that in mice treated with estrogen and estrogen combined with control IgG or Tf antibody was ∼23, 20 and 26 sec, respectively. The same tendency was also observed for PT (Figure 5E). Furthermore, the plasma recalcification time was also accelerated, which was in consistent with the elevated concentration of Tf (Figures 5F and 5G).The shortened APTT, PT and plasma recalcification time might thus contribute to a shortened tail-bleeding time in mice after treatment with estrogen (Figure 5H). As illustrated in Figure 5I, decreased blood flow was observed in the carotid artery of estrogen administration mice by comparison with the control mice. Anti-Tf antibody-treatment reversed the reduction of blood flow and control IgG showed no effect on the reduction (Figure 5I). Estrogen itself showed no effects on coagulant factors (Figures S1A and S1B) and coagulation (Figures S1C and S1D). Directly intravenous administration of estrogen (6 ng/kg) for 10 min (short-term administration) rapidly increased mouse plasma estrogen concentration (Figure S2A), while the Tf concentration (Figure S2B), blood coagulation time (APTT and PT) (Figure S2C), tail bleeding time (Figure S2D) and carotid artery blood flow induced by FeCl_3_ (Figure S3) showed little difference. Therefore our results indicate that estrogen does not directly affect blood coagulation and its effect on hypercoagulability is due to Tf up-regulation.

**Figure 5.**
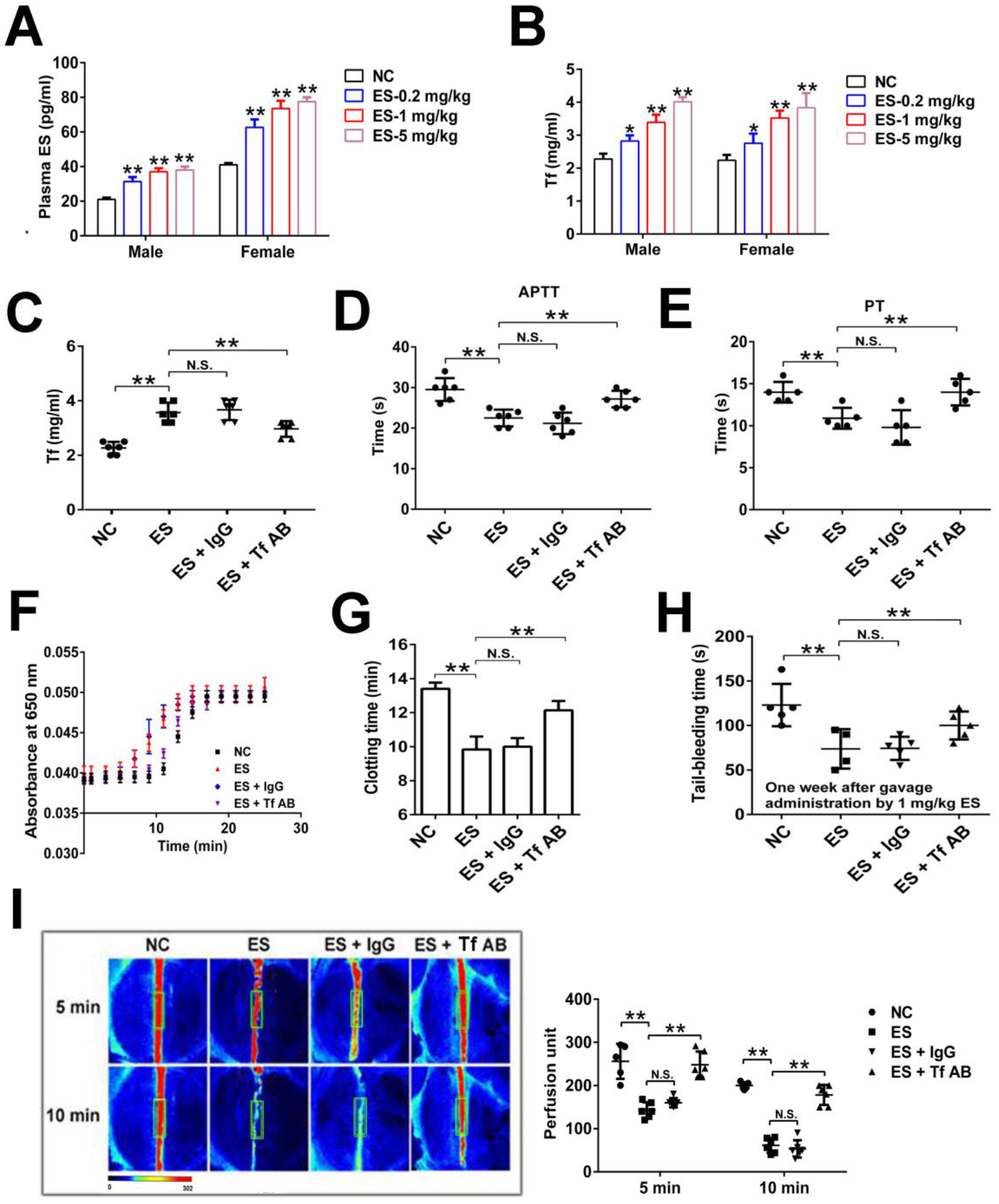
Estrogen induces hypercoagulability and thrombus formation by up-regulating transferrin in mice. The concentration of estrogen **(A)** and Tf **(B)** in plasma of male or female mice after oral gavage administration of different concentrations of estrogen for one week was determined by ELISA. **(C)** Plasma Tf level in the tested mice (NC: negative control, ES: one-week estrogen administration, ES+Tf AB: one-week estrogen administration in the presence of Tf antibody, ES + IgG: one-week estrogen administration in the presence of control IgG), APTT **(D)**, PT **(E)**, plasma recalcification time (**F** and **G**), tail bleeding time **(H)** of the 4 group’s mice were shown. **(I)** Representative images of carotid artery blood flow (left) in FeCl_3_-treated all groups of mice by laser speckle perfusion imaging, and the region of interest (the rectangle in green) was placed in the carotid artery to quantify blood flow change. Relative blood flow in the region of interest was shown (right). Animal experiments were repeated three independent times. Data represent mean ± SD (n = 5-6), \*\**p* < 0.01 by one-way ANOVA with Dunnett’s post hoc test. N.S.: no significance.

### ID, estrogen administration or Tf over-expression aggravates IS which alleviated by Tf interferences

To further investigate the role of Tf *in vivo*, the transient middle cerebral artery occlusion (tMCAO) model was used to assess the effects of Tf on IS. In line with the results in the carotid artery thrombosis formation, ID or estrogen induced a significant increase in infarct volumes (Figures 6A and 6B or Figures 6E and 6F) and more serious functional outcomes (Figures 6C and 6D or Figures 6G and 6H). Importantly, anti-Tf antibody-treatment reversed the sympotoms, while control IgG showed no effect on them (Figures 6A-6H). Furthermore, IS was not aggravated by directly intravenous administration of estrogen (short-term administration) in mice (Figure S4) without up-regulating plasma Tf (Figure S4B).

**Figure 6.**
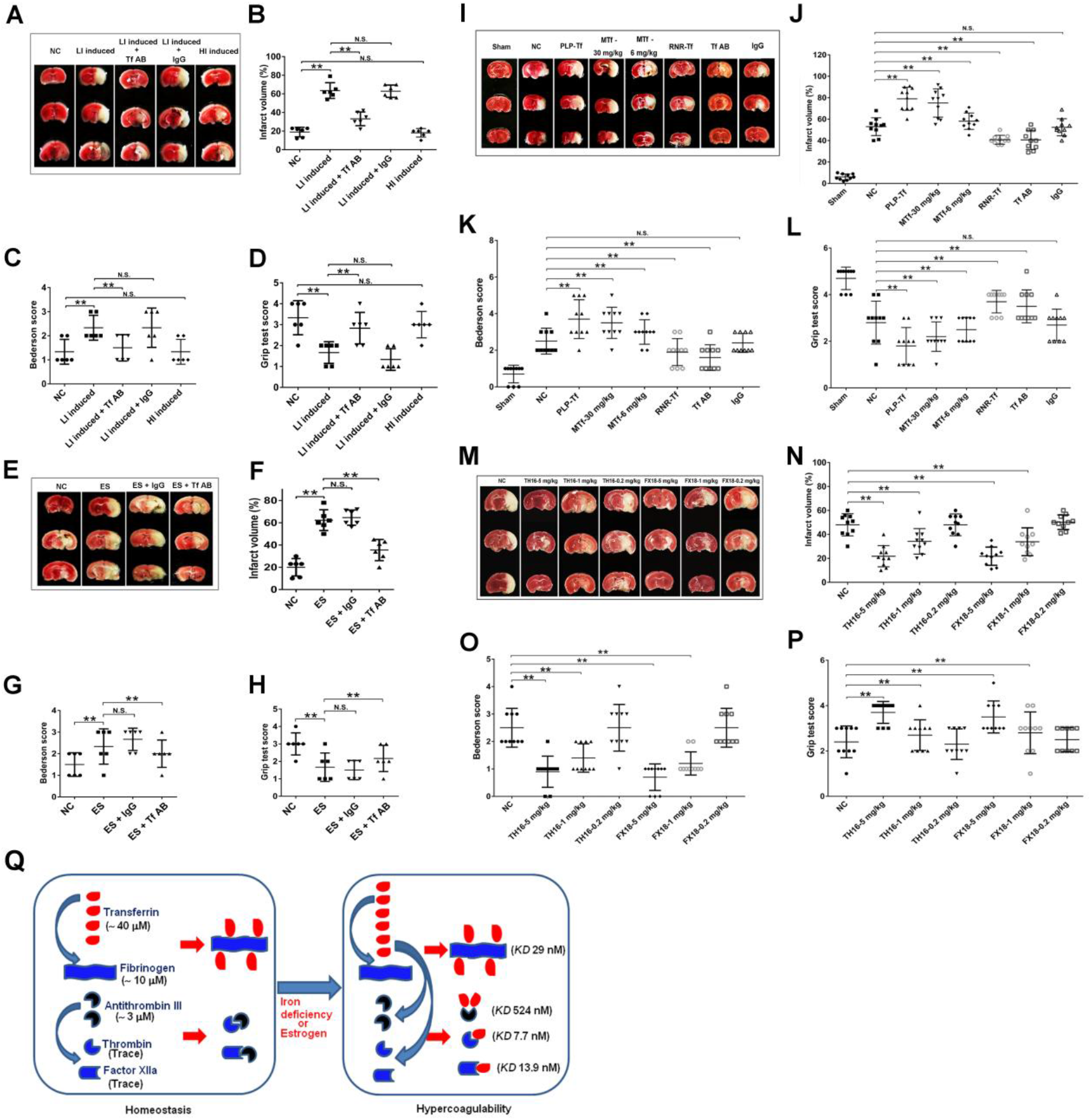
Iron-deficiency, estrogen administration or transferrin over-expression aggravates ischemic stroke. Representative images of 2, 3, 5-triphenyltetrazolium chloride (TTC) stained coronal brain sections **(A)** and quantitative analysis of stained area **(B)** from the 5 groups’ mice as mentioned in “Figure 3” on day 1 after tMCAO. The ischemic infarctions appear in white color and brain infarct volumes as measured by planimetry (% of the whole volume). Bederson score **(C)** and grip test score **(D)** in respective groups were also measured. Representative images of TTC stained coronal brain sections **(E)** and quantitative analysis of stained area **(F)** from the 4 groups’ mice as mentioned in “Figure 5” on day 1 after tMCAO. Bederson score **(G)** and grip test score **(H)** in respective groups were measured. **(I)** Representative TTC stains of the corresponding coronal brain sections from sham-operated (sham), saline (NC), Tf-over-expressed (PLP-Tf), mouse Tf injection (MTf, 6 and 30 mg/kg), Tf knock-down (RNR-Tf), Tf antibody-treated (Tf AB), and control IgG-treated (IgG) mice on day 1 after tMCAO and quantitative analysis of stained area **(J)**. Bederson score **(K)** and grip test score **(L)** in respective groups on day 1 after tMCAO. **(M)** Representative TTC stains of 7 corresponding coronal brain sections from saline- (0.9% sodium chloride, NC), TH16- and FX18-treated (0.2, 1, 5 mg/kg, respectively) mice on day 1 after tMCAO. The two peptides and saline were injected 10 minutes before reperfusion. Brain infarct volumes **(N)**, Bederson score **(O)** and grip test score **(P)** in the respective group on day 1 after tMCAO. **(Q)** The graphical representation of Tf’s central role in coagulation balance and its abnormal up-regulation by ID or estrogen to cause hypercoagulability. Normally, Tf is sequestered by binding with fibrinogen with a molar rate of 4:1 and a high affinity (*KD* ∼29 nM) to keep coagulation balance. Extra Tf induced by ID or estrogen causes hypercoagulability by: 1) binding to AT-III with a molar rate of 2:1 and a low affinity (*KD* ∼524 nM) and blocking inactivation of AT-III towards thrombin and FXa, and 2) interacting with and potentiating thrombin and FXIIa with a molar rate of 1:1 and a high affinity (*KD* ∼ 7.7-13.9 nM). Animal experiments were repeated three independent times. Data represent mean ± SD (n = 6-10), \*\**p* < 0.01 by one-way ANOVA with Dunnett’s post hoc test. N.S.= no significance.

The sequelae of Tf over-expression and knock-down were evaluated. The Tf expression levels were validated by both quantitative real-time polymerase chain reaction (qRT-PCR) and western blot (Figures S5A-S5B). The mice plasma level of Tf was up-regulated or down-regulated by 96 h infection of lentivirus (with Tf over-expression plasmid (PLP-Tf)) or retrovirus (with Tf knock-down plasmid (RNR-Tf)) compared with controls (NC) (Figure S5C). Tf over-expression or administration of Tf induced a significant increase in infarct volumes (Figures 6I and 6J). Furthermore, the increased infarct volumes in Tf over-expressed and Tf-injected mice also translated into more serious functional outcomes (Figures 6K and 6L). Conversely, Tf knock-down or administration of Tf antibody evidently developed smaller infarct volumes (Figures 6I and 6J) and showed better Bederson (Figure 6K) and grip test scores (Figure 6L). Control IgG administration mice showed the similar IS symptom with control mice.

Anti-IS functions of both designed peptides TH16 and FX18, which interfere the interaction between Tf and thrombin/FXIIa, were also tested. Administration of TH16 and FX18 induced a dose-dependently progressive decrease in infarct volumes in mouse IS model (Figures 6M and 6N). Furthermore, the two peptides-treated mice likewise developed improved Bederson (Figure 6O) and grip test scores (Figure 6P). Together, these results indicate that Tf aggravates IS while interference with Tf alleviates it.

## Discussion

Our previous work reported in the accompanying paper has demonstrated that Tf plays a central role to maintain coagulation balance by interacting with coagulation and anti-coagulation factors and suggested that factors up-regulating Tf causes hypercoagulability and thromboembolic diseases. Although recent decades have witnessed a rapidly growing awareness of the association of ID/IDA or CC and thrombosis (Akins et al., 1996; Batur Caglayan et al., 2013; Belman et al., 1990; Benedict et al., 2004; Carr and Ory, 1997; Chasan-Taber and Stampfer, 1998; Coutinho et al., 2015; Devuyst et al., 2000; Farmer et al., 1997; Gialeraki et al., 2016; Hartfield et al., 1997; Jick et al., 2006; Kinoshita et al., 2006; Maguire et al., 2007; Makhija et al., 2008; Nagai et al., 2005), the details of the mechanism remain unclear. Here we show that ID or CC upregulates Tf to induce hypercoagulability and thromboembolic diseases.

The mechanisms of adaptation to ID are associated with suppression of the hepatic hormone hepcidin, which develops consequently to decreased levels of serum ferritin and Tf saturation (Camaschella, 2015). It is suggested that the decreased Tf saturation is partly due to the increased level of Tf, since reduced levels of iron also trigger increases in the synthesis of Tf (Camaschella, 2015). In iron deficiency, the rate of Tf synthesis in the liver increases significantly (2- to 4-fold) (Idzerda et al., 1986). Two adjacent hypoxia response elements (GAAATACGTGCGCTTGTGTGTACGTGCA) containing two the critical (G/A)CGTG core sequences have been identified within the Tf gene enhancer, which are binding sites of HIF-1 (Mukhopadhyay et al., 2000; Rolfs et al., 1997). It has been reported that HIF-1 confers low oxygen up-regulation of Tf gene expression by binding to the two binding sites of HIF-1 present in the Tf enhancer (Rolfs et al., 1997). Previous report has also demonstrated the requirement for HIF-1 in activation of ceruloplasmin transcription by ID (Mukhopadhyay et al., 2000). It is likely that ID up-regulates Tf through HIF-1. Indeed, our result showed that the plasma level of Tf was significantly increased in IDA and IS patients with IDA history, as well as in low-iron diet induced mice (Figures 1A-1C and Figure 3A). This adaptive mechanism may aim to facilitate the absorption of iron or maintain homeostasis. However, an enhanced thrombosis potential may be thus induced because Tf has ability to potentiate thrombin and FXIIa and to block AT-III’s inactivation on coagulation proteases (Figure 6Q). Low-iron diet, not high-iron diet up-regulated Tf, induced hypercoagulability and aggravated IS while anti-Tf antibody reversed the thromboembolic symptoms, suggesting that ID acts as a thrombotic factor by up-regulating Tf.

In chicken, estrogen has been reported to promote the expression of Tf gene in both liver and oviduct (Lee et al., 1978). Elevated Tf was observed in mice treated by estrogen gavage administration for one week as well as in woman VTE patients, who are CC users (Figures 5A and 5B and Figures 1A-1C). We found that estrogen up-regulated Tf expression through EREs located at human Tf gene promoter region and AKT activation (Figure 4) and subsequently induced hypercoagulability (Figures 5C-5H). The ability to induce pro-coagulant state by estrogen was further confirmed by two thrombotic mouse models including carotid artery thrombosis and IS (Figure 5I and Figures 6E-6H). The two diseases were aggravated by estrogen administration while antibody of Tf reversed the disease development.

Two designed peptides (TH16 and FX18), which block Tf-thrombin/FXIIa interaction to inhibit Tf’s potentiation on enzymatic activity of thrombin and FXIIa, inhibited disease development of carotid artery thrombosis and IS induced by either low-iron diet or estrogen application (Figures 6M-6P), confirming further that the two acquired conditions (ID and CC) have the same pathological mechanism to induce thromboembolic development (Figure 6Q). Collectively, the study reveals a mechanistic association between ID, CC, Tf and thrombosis and that factors up-regulating Tf are risk factors of thromboembolic diseases. The central role of Tf to maintain coagulation balance is further confirmed (Figure 6Q). It also provides a novel avenue for the development of anti-thromboembolic medicine associated with acquired conditions such as ID, CC and infection by targeting Tf or interfering Tf-thrombin/FXIIa interaction.

## Methods

### Collection of human samples

According to the clinical protocol approved by the Institutional Review Board of Kunming Institute of Zoology and the Third People’s Hospital of Kunming (Protocol No. KMSYLL-20150101), all human specimens were collected with the informed consent of the patients. IS patients were diagnosed by head computed tomography (CT) and magnetic resonance imaging (MRI) after exhibiting some clinical features (such as inability to move or feel, problems understanding and speaking, blurring of vision or vision loss) (Table S1). All IS patients recruited have IDA history before IS attack. IDA patients were diagnosed by main clinical features, such as low value of red blood cell (RBC), hemoglobin (HGB), hematocrit (HCT), mean corpusular volume (MCV), mean corpusular hemoglobin (MCH) (Table S1). The control group consisted of prospectively recruited healthy volunteers, with matched age whenever possible. Human plasma of the IDA group (n = 454), IS group (n = 453), and healthy volunteers (n = 341) were collected (Table S1). Human CSF of IS group (n = 40) and healthy group (n = 40), as well as thrombus samples from acute IS patients undergoing mechanical thrombectomy were also collected (n=5) (Table S2). Human plasma of VTE patients using CC (n = 30, patients were diagnosed by CT) and healthy volunteers (n = 30, age 30-80) were also collected (Table S3). Immediately following the blood drawing (1.5 % EDTA-Na2 was utilized as an anticoagulant agent), plasma was obtained by centrifuging at 3000 rpm/min for 20 min at 4°C, and then immediately stored at −80°C after being sub-packed.

### Animals and ethics statement

All animal experiments were approved by the Animal Care and Use Committee at Kunming Institute of Zoology (SMKX-2016013) and conformed to the US National Institutes of Health’s Guide for the Care and Use of Laboratory Animals (The National Academies Press, 8th Edition, 2011). C57BL/6J mice (female or male, 8 weeks old) were purchased from Vitalriver Experiment Animal Company (Beijing, China) and housed in a pathogen-free environment.

### Tf concentration determination

Tf concentration in human IS, IDA or VTE patients’ plasma or CSF was determined by ELISA using human Tf ELISA kit (EK12012-96T, Multi Sciences, China) according to the manufacturer’s instructions. Western blot was utilized to determine Tf level in the IS, IDA or VTE patients’ plasma or CSF. The plasma and CSF of patients and healthy controls were separated by 12% sodium dodecyl sulfate-polyacrylamide gel electrophoresis (SDS-PAGE), and then transferred to polyvinylidene difluoride (PVDF) membranes. The membrane was blocked with TBST buffer (50 mM Tris, 150 mM NaCl, 0.1% Tween-20) containing 5% bovine serum albumin at room temperature for 2 h. After washing three times by TBST, the PVDF membrane was incubated the primary antibodies against Tf (1:2000, 11019-RP02, Sino Biological Inc., China) at 4°C overnight, followed by the incubation with the secondary antibody for 1 h at room temperature. After washing with TBST, the membrane was developed with an enhanced chemiluminescence kit (PA112, Tiangen, China) by ImageQuant LAS 4000 mini (GE Healthcare, U.S.A.). To control for plasma or CSF loading and transfer, membranes were stained by Red Ponceau.

### IDA mice induction

C57BL/6J mice (male, 8 weeks old) were used in this experiment. Some mice were subjected to a low-iron diet (Fe < 10 mg/kg) or high-iron diet (Fe > 90 mg/kg), while some mice fed a low-iron diet were injected with a Tf-depleting antibody or control IgG (Tf antibody or control IgG was injected two times/week through the tail vein from the beginning of induction by low-iron forage, and 50 μg per time injection). After 4 weeks induction, blood was collected (1.5 % EDTA-Na2 was utilized as an anticoagulant agent) and then subjected to routine blood tests including RBC, HGB, HCT, MCV, and MCH by using a blood routine test machine (BC-2800Vet, Mindray, China). The blood was also collected (3.8 % sodium citrate was utilized as an anticoagulant agent), and plasma was obtained by centrifuging at 3000 rpm for 30 min at 4°C, and was then used for plasma recalcification time, APTT and PT assays. The tested mice groups were also subjected to test the tail-bleeding time and FeCl_3_-induced carotid artery injury model assays.

### Estrogen administration of mice

C57BL/6J mice (female or male, 8 weeks old) were administrated by oral gavage of estrogen (0.2, 1, and 5 mg/kg, dissolved in 0.9% NaCl) 2 times per week for 1 week. After 1-week induction, plasma of different groups’ mice was extracted and the concentrations of Tf and estrogen were tested by mouse Tf ELISA kit as described above and estrogen ELISA kit (LH-E10064MU, Liuhe, China), respectively.

C57BL/6J mice (female, 8 weeks old) were administrated by gavage of estrogen (1 mg/kg, dissolved in 0.9% NaCl) for two times in a week in the absence or presence of an anti-Tf antibody (Tf AB, Tf antibody was injected for 2 times in a week though tail vein, 50 μg per time injection) or a control IgG. One week after gavage administration (Long-term administration), the mice were also used for plasma recalcification time, APTT, PT, tail-bleeding time and FeCl_3_-induced carotid artery injury model assays. Effect of short-term estrogen administration on coagulation was also evaluated by APTT, PT, tail-bleeding time and FeCl_3_-induced carotid artery injury model assays as the method described in mice intravenously injected by estrogen (6 ng/kg, dissolved by 0.01% DMSO) for 10 min.

### Stroke mouse model

The tMCAO model was applied to induce focal cerebral ischemia as described (Gob et al., 2015; Kraft et al., 2010). Mice were first anesthetized with 2.0 % isoflurane, and tied on a heat controlled operating table (Harvard Apparatus, U.S.A.) to maintain at 37°C during the whole period of surgery. Following a midline skin incision in the neck, the proximal common carotid artery and the external carotid artery (ECA) were ligated, and a standardized silicon rubber-coatednylon monofilament (6023910PK10, Doccol, U.S.A.) was inserted and advanced via the right internal carotid artery to occlude the origin of the right middle cerebral artery. Sixty minutes later, mice were re-anesthetized and the occluding filament was removed to allow reperfusion. Test samples including mouse transferrin (MTf), TH16 and FX18 were injected 10 min before the reperfusion. For determination of ischemic brain volume, mice were sacrificed after tMCAO induction for 24 h, and brains were quickly removed and cut into 2-mm thick coronal sections using a mouse brain slice matrix (Harvard Apparatus, USA). The brain sections were then stained with 2% 2, 3, 5-triphenyltetrazolium chloride (TTC, Sigma, USA). The Bederson score was used to monitor neurological function and the grip test score was used to monitor motor function and coordination.

### Statistical analysis

The data obtained from independent experiments are presented as the mean ± SD. All statistical analyses were two-tailed and with 95% confidence intervals (CI). The results were analyzed using one-way ANOVA with Dunnett’s post hoc test or unpaired t-test by Prism 6 (Graphpad Software) and SPSS (SPSS Inc, USA). Differences were considered significant at p < 0.05.

## Acknowledgments

We thank Dr. Lin Zeng for technical advice and assistance on MS/MS spectra analysis. This work was supported by funding from Chinese Academy of Sciences (XDB31000000, KFJ-BRP-008 and QYZDJ-SSW-SMC012), National Science Foundation of China (331372208), and Yunnan Province (2015HA023) to R.L. and from the National Science Foundation of China (31640071, 31770835 and 81770464), Chinese Academy of Sciences (XDA12020334 and Youth Innovation Promotion Association).

## Author Contributions

X.T., M.F., R.C., Z.Z., C.S., Y.H., J.M., M.X., and C.L. performed the experiments including enzyme activity tests, animal experiments and data analysis; Y.W., Y.D., Y.L., Z.S., and L.Z. collected specimens of IS, IDA and VTE patients; R.L., X.D., and Q.L. prepared the manuscript; R.L. conceived and supervised the project. All authors contributed to the discussions.

## Supplementary Material

Supplementary material includes full methods, Figure S1-S5 and Table S1-S4.

