## Supplemental file for "Iron-deficiency and estrogen are associated with ischemic stroke by up-regulating transferrin to induce hypercoagulability" (from another version of this preprint: version 1, 2019-05-22)

**Materials and Methods**

**Crosslinking assay**

Bis (sulfosuccinimidyl) suberate (BS^3^, 1 mM, 21586, Thermo, USA) was used to crosslink plasma (1 μl, ischemic stroke (IS) patients, iron-deficiency anemia (IDA) patients, venous thromboembolism (VTE) patients using combined oral contraceptives (CC) and normal controls) in HEPES buffer (20 mM, pH 7.4) for 30 min at room temperature. Tris-HCl (10 mM, pH 7.5) was then used to stop the reaction. The crosslinked plasma samples were subjected to 12% SDS-PAGE separation. A rabbit polyclonal antibody against transferrin (Tf, 1:2000, 11019-RP02, Sino Biological, Inc. China) was used to detect Tf. After stripping the PVDF with the stripping buffer (PH1498, Phygene, China), the anti-thrombin (1:2000, ab17199, Abcam, USA) and anti-FXII (1:2000, SC-66752, Santa Cruz, USA) antibody were used to detect prothrombin and FXII, respectively. To control for plasma loading and transfer, membranes were stained by Red Ponceau after transferring.

**Confocal microscopy analysis**

For immunofluorescent assay, frozen slices of thrombus from acute IS patients were blocked for 1 h at room temperature with 1% BSA mixed with 0.2% Triton X-100. To visualize Tf-thrombin/FXIIa puncta, frozen slices were incubated with rabbit polyclonal antibodies against Tf (1: 200, 50817-RP02, Sino Biological Inc, China), mouse polyclonal antibodies against thrombin (1: 200, ab17199, Abcam, UK) or goat polyclonal antibodies against FXII (1: 200, SC-66752, Santa Cruz, USA) at 4 °C overnight, respectively. After washing three times by PBS to remove excess primary antibody, the section was incubated with a fluorescently labeled secondary antibody for 1 h at 37 °C. Cell nuclei were stained with 4, 6-diamidino-2-phenylindole (DAPI, P36941, Lifetechnologies, USA). Images of immunofluorescence were obtained by an Olympus FluoView 1000 confocal microscope according to manufacture’s instructions.

**Effects of iron deficiency (ID) on expression of Tf *in vitro***

A mouse normal liver cell line (BNL CL.2) was used to test effects of ID on expression of Tf *in vitro*. BNL CL.2 was obtained from the Conservation Genetics CAS Kunming Cell Bank and maintained in Dulbecco’s modiﬁed Eagle's medium (DMEM, Gibco Laboratories, USA) containing 10% fetal bovine serum (FBS, MRC, AUS) at 37°C and in the presence of 5% CO_2_. The medium was changed every 3 days and experiments were conducted between the third and eighth passages. Firstly, BNL CL.2 was seeded into 24-well plates at 2.5 × 10^5^ cells per well for 4 h. Iron chelator desferrioxamine (DFO, 0.2, 1 and 5 μM, D9533, Sigma, USA) were then added and incubated with the cells for 24 h to test the amount of Tf by ELISA kit (ab157724, Abcam, USA) and western blot as described. β-actin was used as loading control. The expression of Tf was also tested by quantitative real-time polymerase chain reaction (qRT-PCR) according to the standard procedures. Briefly, cells were harvested. The RNA extraction and cDNA reverse transcription were performed using RNA extraction kit (DP419, Tiangen, China) and reverse transcription kit (A5000, promega, USA) as per the manufacturer’s protocol, respectively. Tf expression was quantified by qRT-PCR (forward primer (5’-3’): GGACGCCATGACTTTGGATG; reverse primer (5’-3’): GCCATGACAGGCACTAGACC for mouse Tf). PCR was performed on the CFX-96 Touch Real-Time Detection System (Bio-Rad, USA).

**Effect of ID on hypoxia-inducible factor-1 (HIF-1)**

BNL CL.2 was seeded into 24-well plates as the method describe above. DFO (0.2, 1 and 5 μM) was then added and incubated with the cells for 8 h to test the HIF-1α by using anti-HIF-1α antibody (1:2000, ab1, Abcam, USA) by western blot as described. β-actin was used as loading control.

**Hypoxia response elements (HRE) assays**

pUC57 plasmid of wild-type human *Tf* gene range from -5500 to +2094 (GenBank with Accession number (NC_000003)) was constructed. Nucleotide +1 is the A base of the ATG-translation initiation codon. Mutant plasmid of two adjacent HRE (ACGTG) within the enhancer region of *Tf* gene was also constructed and all the bases in HRE were mutated to A base. Human hepatocarcinoma HepG2 cells (Conservation Genetics CAS Kunming Cell Bank) were transfected with wild-type or variant HRE reporter of Tf expression plasmid or pUC57 empty plasmid by using lipofectamine 2000. The transfection procedure was performed according to manufacturer's manual. Eight hour after transfection, the cells were cultured in DMEM for another 12 h. Different concentrations of DFO (0.2, 1 and 5 μM) were added and incubated with the cells for 24 h, the cell supernatant was then collected and the concentration of Tf in the supernatant was tested by ELISA kit (EK12012-96T, Multi Sciences, China).

**Effects of estrogen on expression of Tf *in vitro***

BNL CL.2 was used to test effects of estrogen on expression of Tf *in vitro*. Firstly, BNL CL.2 was seeded into 24-well plates at 2.5 × 10^5^ cells per well for 4 h as the method describe above. Estrogen (Estradiol, E8875, Sigma, USA) (0.1, 1, and 10 ng/ml, dissolved in 0.01% Dimethyl sulfoxide (DMSO)) were then added and incubated with the cells for 24 h to test the amount of Tf by ELISA kit (ab157724, Abcam, USA) and western blot as described. β-actin was used as loading control.

**Estrogen response elements (EREs) assays**

pUC57 plasmid of wild-type mouse *Tf* gene range from -3070 to +2094 (GenBank with Accession number ([22041](https://www.ncbi.nlm.nih.gov/sites/entrez?db=gene&cmd=Retrieve&dopt=full_report&list_uids=22041))) was also constructed. Nucleotide +1 is the A base of the ATG-translation initiation codon. Mutant plasmid of estrogen-responsive promoter region (EREs) between -1021 and -1009 (GGAGAGGATGGCC) of *Tf* gene was also constructed and all the bases in EREs region (GGAGAGGATGGCC) were mutated to A base. BNL CL.2 cells were transfected with wild-type or variant EREs reporter of Tf expression plasmid or pUC57 empty plasmid by using lipofectamine 2000. The transfection procedure was performed according to manufacturer's manual. Eight hour after transfection, the cells were cultured in DMEM for another 12 h. Different concentrations of estrogen (0.1, 1, and 10 ng/ml, dissolved in 0.01% DMSO) were added and incubated with the cells for 24 h, the cell supernatant was then collected and the concentration of Tf in the supernatant was tested by ELISA kit as described above.

**Effects of estrogen on transcriptional factor C/EBPα**

BNL CL.2 cells were used to test the effects of estrogen on transcriptional factor C/EBPα. BNL CL.2 was first seeded into 24-well plates at 2.5 × 10^5^ cells per well for 4 h as the method describe above. Different concentration of estrogen (0.1, 1, and 10 ng/ml, dissolved in 0.01% DMSO) was incubated with the cells in the absence or presence of wortmannin (10 nM, MB5799-5MG, Dalianmeilun, China) for 2 h to test phosphorylation of protein kinase B (AKT), glycogen synthase kinase-3β (GSK3β), and CCAAT/enhancer binding protein-α (C/EBPα) by western blot using rabbit polyclonal anti-[phospho-AKT](https://www.cellsignal.com/products/primary-antibodies/phospho-akt-ser473-d9e-xp-rabbit-mab/4060?N=4294956287&Ntt=akt&fromPage=plp) antibody (1:5000, 9474, CST, USA), rabbit polyclonal anti-phospho-GSK3β antibody (1:2000, AF2016, Affinity, USA), and rabbit polyclonal anti-[phospho-C/EBPα](https://www.cellsignal.com/products/primary-antibodies/phospho-akt-ser473-d9e-xp-rabbit-mab/4060?N=4294956287&Ntt=akt&fromPage=plp) antibody (1:2000, 2844, CST, USA), respectively as the method described above. To test C/EBPα expression induced by estrogen, BNL CL.2 was first seeded into 24-well plates at 2.5 × 10^5^ cells per well for 4 h, different concentration of estrogen (0.1, 1, 10 ng/ml) was then added in the absence or presence of wortmannin (10 nM) and incubated with the cells for another 24 h. A rabbit polyclonal anti-C/EBPα antibody (1:5000, 2843, CST, USA) was used to test the expression of C/EBPα.

**Effects of estrogen on coagulation factors and and plasma recalcification time**

Effects of estrogen on coagulation factors (FXIIa and thrombin) were tested by using corresponding chromogenic substrates. The testing enzyme was incubated with different concentrations of estrogen (0.1, 1, and 10 ng/ml, dissolved in 0.01% DMSO) in 60 μl Tris buffer (50 mM, pH 7.4) for 5 min, and a certain concentration of chromogenic substrate was then added. The absorbance at 405 nm was monitored immediately and the kinetic curve was recorded by using an enzyme-labeled instrument (Epoch, BioTek, U.S.A.) for 30 min. Human α-Thrombin (T6884, Sigma, USA, 20 nM) and human factor α-XIIa (HFXIIa 1212a, Enzyme Research Laboratories, USA, 20 nM) were reacted with 0.2 mM chromogenic substrate of *H-D*-Phe-Pip-Arg-*p*Na·2HCl (CS-01, [Hyphen Biomed](http://www.baidu.com/link?url=TxVDu_bCpdBR3DTeXrmLAh9nmLYo5YCUZgy9MwnrAVq37hHB93oK3MYQ2H67XVqnno8KJMPiLczgL8OvV_aDewKQsHRflYHBaDUN4CdYmO3), France) and *H-D*-Pro-Phe-Arg-*p*NA·2HCl (CS-31, [Hyphen Biomed](http://www.baidu.com/link?url=TxVDu_bCpdBR3DTeXrmLAh9nmLYo5YCUZgy9MwnrAVq37hHB93oK3MYQ2H67XVqnno8KJMPiLczgL8OvV_aDewKQsHRflYHBaDUN4CdYmO3), France), respectively.

To test plasma recalcification time, 20 μl of plasma was incubated with different concentrations of estrogen (0.1, 1, 10 ng/ml, dissolved in 0.01% DMSO) in 60 μl of HEPES buffer (20 mM HEPES, 150 mM NaCl, pH 7.4) for 10 min at 37°C, 60 μl of CaCl_2_ (25 mM) was then added and incubated for 10 min at 37°C. The clotting was monitored at 650 nm, and the clotting time was calculated by measuring the time to half maximal increase in absorbance.

**APTT and PT assays**

For APTT assay, 50 μl of APTT reagent (F008-1, Nanjing Jiancheng Bioengineering Institute, China) was incubated with 50 μl plasma for 3 min at 37°C, and then 50 μl of CaCl_2_ (25 mM) preheated at 37°C was added to test the clotting time by monitoring the absorbance at 405 nm using a semi-automatic coagulation analyzer (ThromboScreen 400c, pacific hemostasis, USA). To test PT, 50 μl plasma preheated at 37°C was mixed with 100 μl PT reagent (F007, Nanjing Jiancheng Bioengineering Institute, China) preheated at 37°C for 15 min by monitoring the absorbance at 405 nm.

**Bleeding time measurement**

Briefly, the mouse tail was cut 2 mm from the tip, and was carefully immersed in 20 ml of saline warmed at 37°C. Bleeding time was recorded for the stream of blood to cease.

**FeCl_3_-induced carotid artery injury model**

Briefly, the mice were firstly anaesthetized with 2.0% isoflurane, and core body temperature was maintained at 37°C throughout surgery. One of the carotid arteries was exposed by cervical incision, and was then separated from the adherent tissue and vagus nerve. Thrombosis was induced by applying a piece (2 × 2 mm) of filter paper that was pre-soaked with 10% (w/v) FeCl_3_ solution to the exposed mice carotid artery. Blood flow in the carotid artery of all groups was measured by laser speckle perfusion imaging (PeriCam PSI, HR, Sweden) at 5 min and 10 min after FeCl_3_ induction, respectively, and the perfusion unit of region of interest (ROI) was also recorded.

**Generation of lentiviral and retroviral vectors and virus package for Tf over-expression or knockdown**

The coding region of mouse Tf (GenBank: BC092046.1) was synthesized and cloned into pLVX-Puro lentiviral plasmid (Clontech, USA) between EcoRI and BamHI sites. The empty pLVX-Puro plasmid was used as blank control. The oligonucleotides of shRNA sequence targeting Tf (sh-Tf) and its scrambled control shRNA (sh-scr) sequence were synthesized by Sangon Biotech (Shanghai, China). The sequence of sh-Tf is 5’-AAGAATCTGAAGCAGGAAGAC TTCAAGAGAGTCTTCCTGCTTCAGATTCTTCTTTTTT-3’ (Forward oligo), while that for sh-scr is 5’- ACTGAGACTGAACAAAGGAAGTTCAAGAGACTTCCTTTGTTCAGTCTCAGTCTTTTTT-3’ (Forward oligo). The oligonucleotides of sh-Tf and sh-scr including the digested sites of BamHI and EcoRI were annealed with their complementary sequences and inserted into BamHI and EcoRI site of the RNAi-Ready pSIREN-RetroQ retroviral vector (Clontech, USA) to form retroviral vectors of psh-Tf and psh-scr, respectively. Lentiviral vector for Tf over-expression and retroviral vector for Tf knockdown were transfected into murine embryonic liver cells (BNL CL.2 cells, Conservation Genetics CAS Kunming cell Bank, China) to test the efficiency of Tf over-expression or knockdown using western blot and qRT-PCR. The primers for PCR were: forward primer (5’-3’): GGACGCCATGACTTTGGATG and reverse primer (5’-3’): GCCATGACAGGCACTAGACC). The transfection procedure was performed according to the lipofectamine 2000 (11668019, Life technologies, USA) manufacturer's instructions.

HEK 293T cells (Conservation Genetics CAS Kunming cell Bank, China) and EcoPack™ 2–293 cells (Clontech, USA) were used to package lentiviruses and retroviruses, respectively. For each virus preparation, 1.5 × 10^7^ HEK 293T cells or EcoPack™ 2–293 cells were seeded in a T75 flask for 24 h before transfection. pLVX-Puro empty lentiviral vector (20 μg) or with mouse Tf gene and the packaging plasmids including psPAX2 (15 μg, Addgene Inc., USA) and pMD2G (10 μg, Addgene Inc., USA) were co-transfected into HEK 293T cells by lipofectamine 2000. Each flask of EcoPack™2–293 cells was transfected with 25 μg retroviral vector psh-Tf or psh-scr in 70 μl lipofectamine 2000. The medium was changed with fresh medium after 8 h transfection, and the supernatants containing viruses were harvested after 48 h from the first transfection and concentrated by ultracentrifugation using a P28S rotor (Himac, Japan) at 25,000 rpm for 3 h at 4°C, and then immediately stored at -80°C after being sub-packed. HEK 293T cells (100 μl, 4 × 10^4^ cells) were seeded in a 96-well plate to measure virus titer. Briefly, after the infection by six gradients of (dilution ratio: 10, 10^2^, 10^3^, 10^4^, 10^5^, and 10^6^) lentiviruses (pLVX-Puro-Tf, and negative control) or retroviruses (psh-Tf, and psh-scr) for 48 h, puromycin (2 μg/ml, ab141841, Abcam, USA) was added to screen infected cells for 48 h and virus titer was measured by infection cells (living cells) counting.

**Anti-Tf polyclonal antibody preparation**

Mouse Tf (T0523, Sigma, USA) was used to immunize rabbit to produce anti-Tf polyclonal antibody. Briefly, rabbits (male, 2 kg) were primed by subcutaneous injections of 500 μg mouse Tf in 1 ml complete Freund’s adjuvant (Sigma-Aldrich Inc.) on day 0, followed by subcutaneous booster injections of the half dose of the antigens in incomplete Freund’s adjuvant (Sigma-Aldrich Inc.) on day 14, 28, and 42. The titer and specificity of polyclonal antibodies were screened by indirect ELISA. Anti-Tf polyclonal antibodies were then purified from rabbit serum on a protein G column from Amersham Biosciences (Piscataway, NJ).

**Peptides synthesis**

TH16 (RIGKHSRTRYERNIEK), FX18 (RRNHSCEPCQTLAVRSYR) and their scrambled peptides (TH16-scr RKKGIRRYTERHSNIE, FX18-scr SCPTHYSRQRCRNAVLER) were synthesized on an automatic peptide synthesizer (PerSeptive Biosystems, Boston, MA, USA) according to manufacture’s instruction. A method coupling matrix-assisted laser desorption/ionization time-of-flight mass spectrometry and reverse phase high performance liquid chromatography techniques was performed to determine the purity of peptides to be greater than 98%.

**Supplementary Figures and Figure legends**

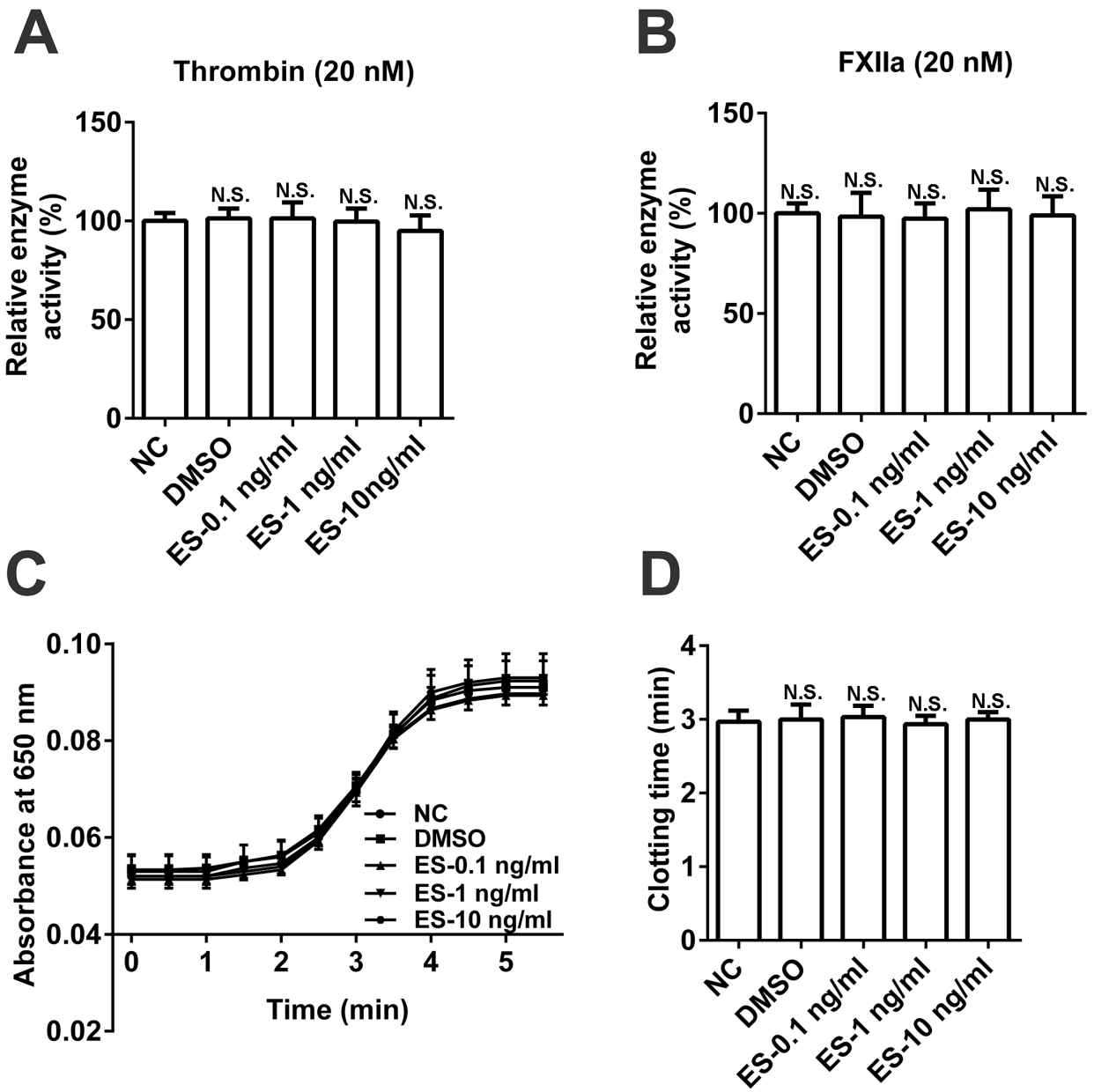

**Figure S1. Effects of estrogen on coagulation factors and blood coagulation.** Effect of estrogen on thrombin **(A)**, FXIIa **(B)**, and plasma recalcification time **(C)**. **(D)** Quantification of **(C)** by measuring the time to half maximal increase in absorbance. ES: estrogen. Data represent mean ± SD of 4 independent experiments. N.S.: no significance.

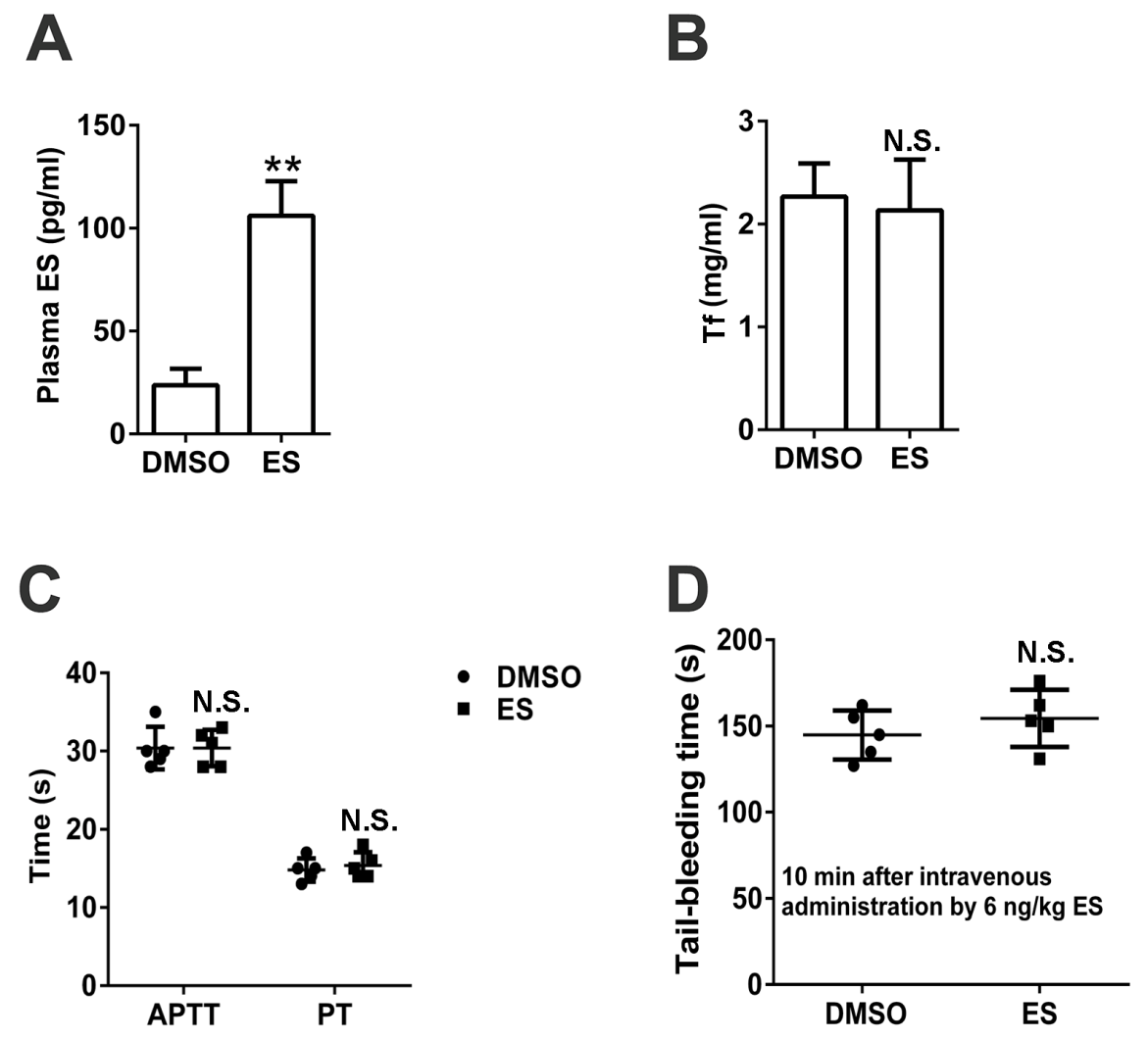

**Figure S2. Effects of short-term estrogen administration on coagulation.** After estrogen (6 ng/kg) or DMSO intravenous injection into C57BL/6J mice for 10 min (n = 10), plasma concentration of estogen **(A)** and Tf **(B)**, APTT and PT **(C)**, and tail bleeding time **(D)** were tested. Data represent mean ± SD (n = 5), ***p* < 0.01 by unpaired t-test. N.S.: no significance.

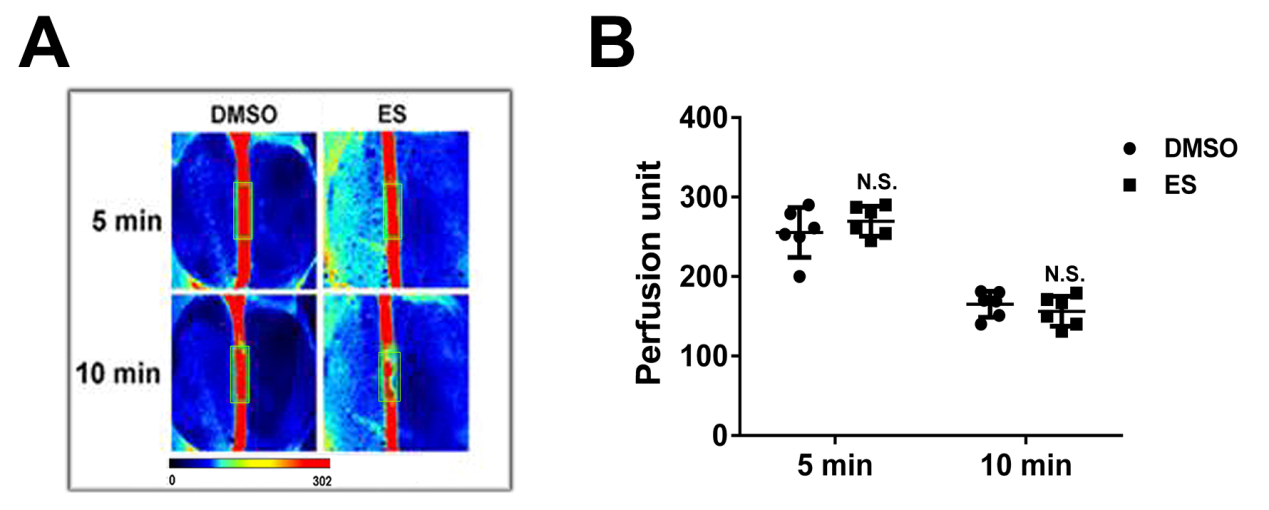

**Figure S3. Effects of short-term estrogen administration on thrombosis.** After estrogen (6 ng/kg) or DMSO intravenous injection into C57BL/6J mice for 10 min, representative images of carotid artery blood flow **(A)**, and quantification of carotid artery blood flow **(B)** were shown. Data represent mean ± SD (n = 6). N.S.: no significance.

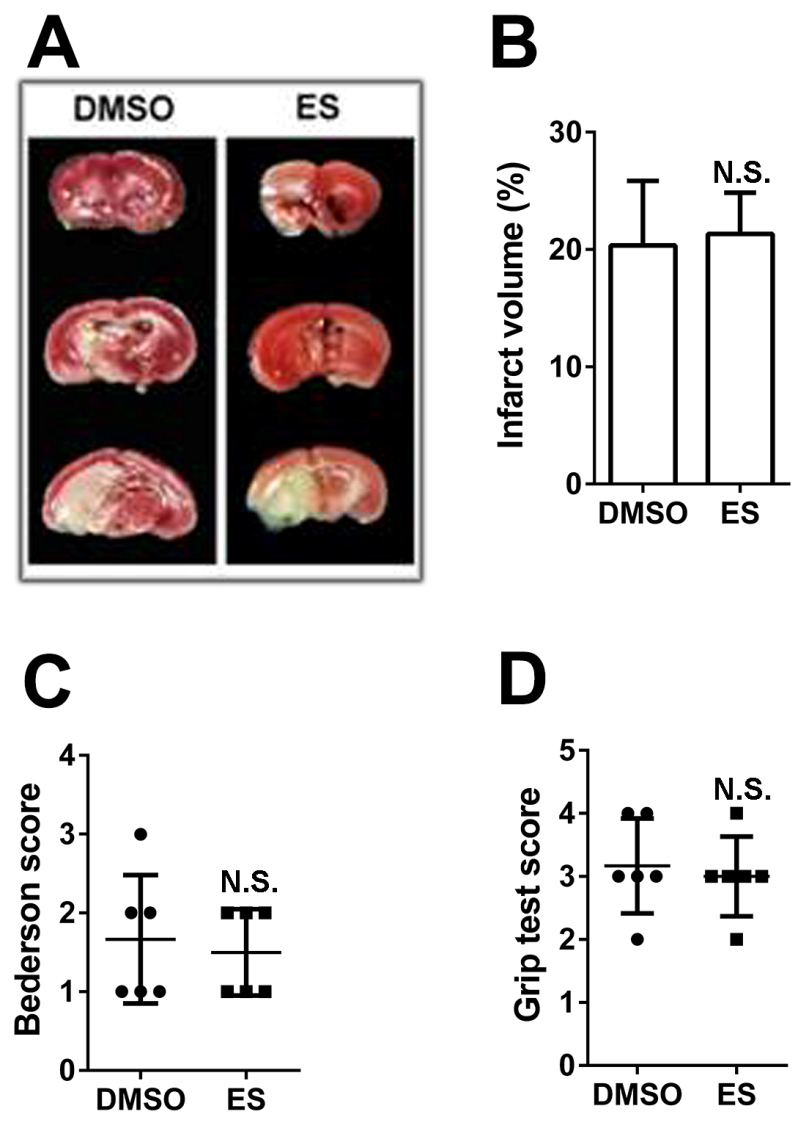

**Figure S4. Effects of short-term estrogen administration on stroke model.** After estrogen (6 ng/kg) or DMSO intravenous injection into C57BL/6J mice for 10 min, representative coronal brain sections stained with TTC from mice on day 1 after tMCAO **(A)**. The ischemic infarctions appear white. Brain infarct volumes **(B)** as measured by planimetry (% of the whole volume), Bederson score **(C)** and grip test score **(D)** in respective group on day 1 after tMCAO were also shown. ES: estrogen. Data represent mean ± SD (n=6). N.S.: no significance.

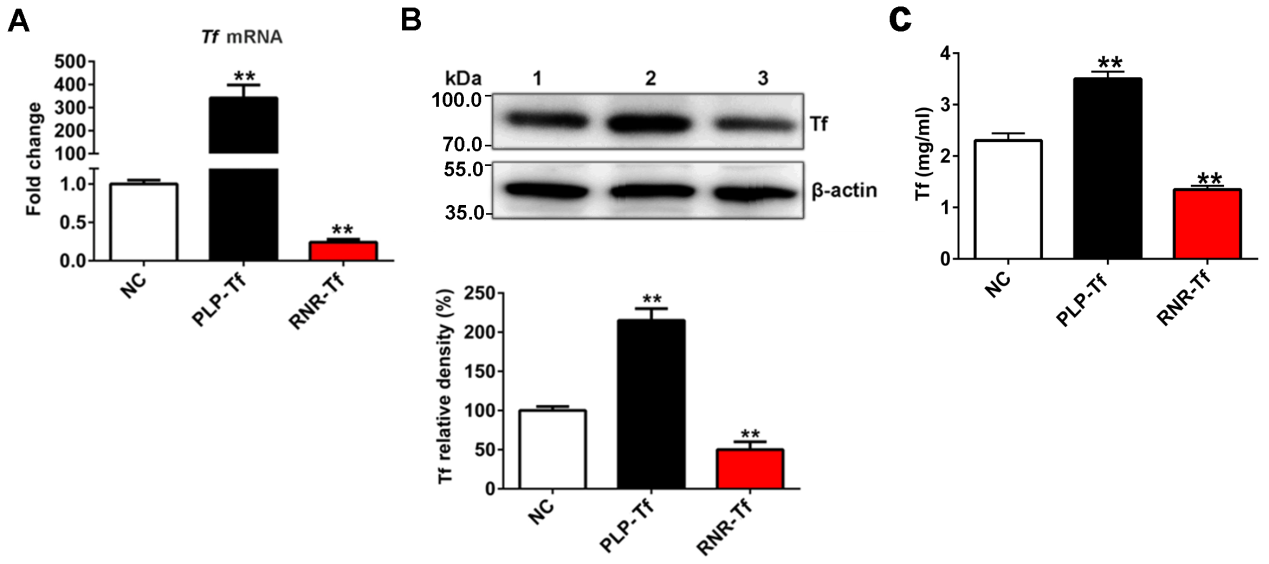

**Figure S5.** **Construction of Tf over-expression and knockdown vectors.** **(A)** Tf mRNA levels of BNL CL.2 cells after transfecting the over-expression or knockdown plasmid of Tf were determined by qRT-PCR. **(B)** Tf levels in the BNL CL.2 cells were determined by western blot (top, Lane 1: control (NC), Lane 2: over-expression (PLP-Tf), Lane 3: knockdown (RNR-Tf)). The quantification of the result of western blot was also shown (bottom). Data represent mean ± SD of six independent experiments, ***p* < 0.01 by one-way ANOVA with Dunnett’s post hoc test. **(C)** The C57BL/6J mice plasma level of Tf after 96 h infection of lentivirus (with Tf over-expression plasmid (PLP-Tf)) or retrovirus (with Tf knock-down plasmid (RNR-Tf)) compared with controls (NC) was determined by ELISA. Data represent mean ± SD (n = 10 ), ***p* < 0.01 by one-way ANOVA with Dunnett’s post hoc test.

**Supplementary Tables**

**Table S1** Clinical features, and plasma Tf concentrations in patients with IS, IDA, and healthy controls.

| IS | | | | IDA | | | | Controls (Normal physical examinees) | | | |
| --- | --- | --- | --- | --- | --- | --- | --- | --- | --- | --- | --- |
| Gender | Age  (years) | Main clinical features | Tf  (mg/ml) | Gender | Age  (years) | Main clinical  features | Tf (mg/ml) | Gender | Age  (years) | Main clinical features | Tf (mg/ml) |
| Male | 69 | IMF; PUS | 5.368 | Female | 60 | Low value of RBC, HGB, HCT, MCV, MCH | 3.120 | Male | 39 | NA | 2.880 |
| Female | 71 | IMF; BVVL; | 4.103 | Male | 66 | Low value of RBC, HGB, HCT, MCV, MCH | 4.044 | Female | 41 | NA | 3.240 |
| Female | 86 | IMF; BVVL; PUS | 3.223 | Female | 51 | Low value of RBC, HGB, HCT, MCV, MCH | 2.184 | Male | 21 | NA | 2.290 |
| Female | 66 | IMF; BVVL; PUS | 3.179 | Female | 74 | Low value of RBC, HGB, HCT, MCV, MCH | 4.476 | Male | 25 | NA | 1.810 |
| Male | 90 | BVVL; PUS | 5.192 | Male | 64 | Low value of RBC, HGB, HCT, MCV, MCH | 2.880 | Female | 39 | NA | 4.250 |
| Female | 90 | IMF; BVVL; PUS | 3.641 | Male | 42 | Low value of RBC, HGB, HCT, MCV, MCH | 3.060 | Female | 32 | NA | 3.720 |
| Female | 38 | IMF; BVVL; PUS | 3.190 | Female | 77 | Low value of RBC, HGB, HCT, MCV, MCH | 2.760 | Female | 35 | NA | 2.630 |
| Male | 47 | IMF; BVVL; PUS | 3.784 | Female | 40 | Low value of RBC, HGB, HCT, MCV, MCH | 2.124 | Male | 23 | NA | 2.860 |
| Female | 80 | IMF; BVVL | 4.411 | Female | 73 | Low value of RBC, HGB, HCT, MCV, MCH | 2.256 | Female | 27 | NA | 2.110 |
| Female | 57 | IMF; BVVL | 5.665 | Female | 74 | Low value of RBC, HGB, HCT, MCV, MCH | 2.640 | Male | 53 | NA | 2.460 |
| Male | 36 | IMF; BVVL; PUS | 4.180 | Female | 66 | Low value of RBC, HGB, HCT, MCV, MCH | 2.952 | Male | 23 | NA | 2.430 |
| Male | 67 | IMF; BVVL; | 4.092 | Male | 70 | Low value of RBC, HGB, HCT, MCV, MCH | 2.268 | Male | 33 | NA | 2.200 |
| Male | 35 | IMF; BVVL; PUS | 4.686 | Male | 73 | Low value of RBC, HGB, HCT, MCV, MCH | 3.432 | Female | 34 | NA | 4.000 |
| Female | 80 | IMF; BVVL; PUS | 3.201 | Male | 72 | Low value of RBC, HGB, HCT, MCV, MCH | 4.092 | Male | 30 | NA | 2.290 |
| Female | 76 | IMF; BVVL; PUS | 2.200 | Female | 50 | Low value of RBC, HGB, HCT, MCV, MCH | 2.460 | Female | 26 | NA | 4.020 |
| Male | 75 | IMF; PUS | 4.125 | Male | 67 | Low value of RBC, HGB, HCT, MCV, MCH | 1.860 | Female | 32 | NA | 2.820 |
| Female | 77 | IMF; BVVL; PUS | 4.488 | Female | 78 | Low value of RBC, HGB, HCT, MCV, MCH | 9.636 | Male | 29 | NA | 4.120 |
| Female | 57 | IMF; BVVL; PUS | 4.950 | Male | 39 | Low value of RBC, HGB, HCT, MCV, MCH | 2.784 | Female | 76 | NA | 2.200 |
| Female | 76 | IMF; BVVL; PUS | 3.520 | Female | 72 | Low value of RBC, HGB, HCT, MCV, MCH | 5.064 | Female | 61 | NA | 2.650 |
| Male | 75 | IMF; BVVL | 8.404 | Female | 78 | Low value of RBC, HGB, HCT, MCV, MCH | 3.408 | Female | 37 | NA | 3.590 |
| Female | 81 | IMF; BVVL; PUS | 5.995 | Female | 84 | Low value of RBC, HGB, HCT, MCV, MCH | 2.772 | Female | 47 | NA | 2.800 |
| Male | 63 | IMF; BVVL; PUS | 2.915 | Male | 77 | Low value of RBC, HGB, HCT, MCV, MCH | 2.964 | Male | 31 | NA | 3.000 |
| Male | 69 | IMF; BVVL; PUS | 4.037 | Male | 72 | Low value of RBC, HGB, HCT, MCV, MCH | 5.244 | Female | 38 | NA | 2.840 |
| Male | 54 | IMF; PUS | 3.960 | Male | 64 | Low value of RBC, HGB, HCT, MCV, MCH | 2.820 | Female | 25 | NA | 3.630 |
| Female | 77 | IMF; BVVL | 6.578 | Male | 70 | Low value of RBC, HGB, HCT, MCV, MCH | 2.880 | Female | 49 | NA | 4.700 |
| Male | 67 | IMF; BVVL; PUS | 5.291 | Male | 66 | Low value of RBC, HGB, HCT, MCV, MCH | 3.348 | Male | 44 | NA | 2.900 |
| Male | 45 | IMF; BVVL; PUS | 3.476 | Male | 68 | Low value of RBC, HGB, HCT, MCV, MCH | 8.832 | Female | 52 | NA | 3.200 |
| Female | 80 | IMF; BVVL; PUS | 5.280 | Female | 48 | Low value of RBC, HGB, HCT, MCV, MCH | 6.120 | Female | 28 | NA | 2.300 |
| Male | 43 | IMF; BVVL; PUS | 4.917 | Female | 63 | Low value of RBC, HGB, HCT, MCV, MCH | 7.044 | Female | 67 | NA | 3.660 |
| Female | 76 | IMF; PUS | 3.773 | Female | 60 | Low value of RBC, HGB, HCT, MCV, MCH | 2.844 | Female | 65 | NA | 2.390 |
| Female | 77 | IMF; BVVL; PUS | 3.531 | Female | 48 | Low value of RBC, HGB, HCT, MCV, MCH | 5.124 | Female | 21 | NA | 2.300 |
| Female | 69 | IMF; BVVL; PUS | 3.762 | Female | 62 | Low value of RBC, HGB, HCT, MCV, MCH | 3.192 | Female | 25 | NA | 3.080 |
| Female | 73 | BVVL; PUS | 3.894 | Female | 45 | Low value of RBC, HGB, HCT, MCV, MCH | 2.964 | Female | 23 | NA | 2.900 |
| Female | 76 | IMF; BVVL; PUS | 5.291 | Female | 48 | Low value of RBC, HGB, HCT, MCV, MCH | 8.328 | Male | 33 | NA | 2.700 |
| Female | 75 | IMF; BVVL; PUS | 3.058 | Female | 63 | Low value of RBC, HGB, HCT, MCV, MCH | 4.968 | Male | 29 | NA | 1.890 |
| Female | 53 | IMF; BVVL | 6.490 | Male | 73 | Low value of RBC, HGB, HCT, MCV, MCH | 4.500 | Male | 69 | NA | 3.100 |
| Female | 70 | IMF; BVVL | 3.289 | Female | 60 | Low value of RBC, HGB, HCT, MCV, MCH | 2.196 | Female | 31 | NA | 2.820 |
| Female | 2 | IMF; BVVL | 4.048 | Female | 63 | Low value of RBC, HGB, HCT, MCV, MCH | 2.652 | Male | 61 | NA | 2.830 |
| Male | 42 | IMF; BVVL; PUS | 5.016 | Female | 66 | Low value of RBC, HGB, HCT, MCV, MCH | 3.624 | Male | 23 | NA | 2.690 |
| Female | 75 | IMF; BVVL; PUS | 5.016 | Female | 68 | Low value of RBC, HGB, HCT, MCV, MCH | 4.464 | Male | 42 | NA | 3.200 |
| Female | 81 | IMF; BVVL; PUS | 3.179 | Female | 83 | Low value of RBC, HGB, HCT, MCV, MCH | 4.440 | Male | 16 | NA | 3.810 |
| Male | 87 | IMF; BVVL; PUS | 2.552 | Male | 59 | Low value of RBC, HGB, HCT, MCV, MCH | 4.596 | Female | 33 | NA | 2.300 |
| Female | 80 | IMF; BVVL; PUS | 4.158 | Male | 83 | Low value of RBC, HGB, HCT, MCV, MCH | 4.440 | Male | 42 | NA | 3.100 |
| Male | 75 | IMF; PUS | 4.895 | Male | 83 | Low value of RBC, HGB, HCT, MCV, MCH | 2.616 | Male | 28 | NA | 2.050 |
| Female | 67 | IMF; BVVL; PUS | 3.718 | Female | 31 | Low value of RBC, HGB, HCT, MCV, MCH | 3.912 | Female | 32 | NA | 3.910 |
| Female | 73 | IMF; PUS | 5.291 | Female | 40 | Low value of RBC, HGB, HCT, MCV, MCH | 4.692 | Female | 38 | NA | 3.600 |
| Male | 69 | IMF; BVVL; PUS | 6.380 | Female | 73 | Low value of RBC, HGB, HCT, MCV, MCH | 4.188 | Male | 41 | NA | 4.140 |
| Female | 71 | IMF; BVVL; PUS | 5.060 | Female | 50 | Low value of RBC, HGB, HCT, MCV, MCH | 2.328 | Male | 44 | NA | 1.470 |
| Male | 51 | IMF; BVVL | 8.316 | Male | 29 | Low value of RBC, HGB, HCT, MCV, MCH | 3.252 | Male | 35 | NA | 3.120 |
| Male | 66 | IMF; BVVL | 4.653 | Male | 60 | Low value of RBC, HGB, HCT, MCV, MCH | 2.904 | Female | 57 | NA | 3.420 |
| Male | 63 | IMF; BVVL; PUS | 6.897 | Male | 28 | Low value of RBC, HGB, HCT, MCV, MCH | 9.012 | Female | 35 | NA | 4.140 |
| Male | 51 | IMF; BVVL; PUS | 5.577 | Female | 73 | Low value of RBC, HGB, HCT, MCV, MCH | 3.696 | Male | 40 | NA | 2.260 |
| Male | 65 | IMF; BVVL; PUS | 6.490 | Female | 69 | Low value of RBC, HGB, HCT, MCV, MCH | 2.724 | Female | 60 | NA | 2.380 |
| Male | 48 | IMF; BVVL; PUS | 5.973 | Female | 68 | Low value of RBC, HGB, HCT, MCV, MCH | 3.360 | Female | 31 | NA | 2.460 |
| Male | 51 | IMF; BVVL; PUS | 8.459 | Male | 70 | Low value of RBC, HGB, HCT, MCV, MCH | 9.300 | Female | 65 | NA | 2.630 |
| Male | 66 | IMF; BVVL; PUS | 6.556 | Male | 50 | Low value of RBC, HGB, HCT, MCV, MCH | 2.268 | Female | 28 | NA | 4.270 |
| Female | 76 | IMF; BVVL; PUS | 6.468 | Female | 69 | Low value of RBC, HGB, HCT, MCV, MCH | 4.116 | Female | 37 | NA | 2.700 |
| Male | 63 | IMF; BVVL; PUS | 6.963 | Male | 68 | Low value of RBC, HGB, HCT, MCV, MCH | 2.592 | Female | 44 | NA | 2.900 |
| Female | 66 | IMF; BVVL; PUS | 5.709 | Female | 74 | Low value of RBC, HGB, HCT, MCV, MCH | 5.304 | Female | 26 | NA | 2.330 |
| Male | 48 | IMF; BVVL | 7.865 | Male | 34 | Low value of RBC, HGB, HCT, MCV, MCH | 2.640 | Female | 51 | NA | 2.410 |
| Male | 66 | IMF; BVVL | 7.040 | Male | 50 | Low value of RBC, HGB, HCT, MCV, MCH | 2.340 | Male | 41 | NA | 3.000 |
| Female | 76 | IMF; BVVL; PUS | 5.445 | Female | 21 | Low value of RBC, HGB, HCT, MCV, MCH | 4.920 | Male | 45 | NA | 2.570 |
| Male | 79 | IMF; BVVL | 6.160 | Male | 55 | Low value of RBC, HGB, HCT, MCV, MCH | 1.992 | Female | 56 | NA | 2.380 |
| Male | 68 | IMF; PUS | 5.038 | Male | 65 | Low value of RBC, HGB, HCT, MCV, MCH | 9.864 | Female | 32 | NA | 3.100 |
| Female | 65 | IMF; BVVL; PUS | 6.897 | Male | 27 | Low value of RBC, HGB, HCT, MCV, MCH | 8.724 | Male | 37 | NA | 3.070 |
| Female | 49 | IMF; BVVL; PUS | 5.357 | Male | 63 | Low value of RBC, HGB, HCT, MCV, MCH | 3.792 | Male | 37 | NA | 3.290 |
| Female | 49 | IMF; BVVL; PUS | 6.996 | Male | 71 | Low value of RBC, HGB, HCT, MCV, MCH | 1.824 | Female | 64 | NA | 4.070 |
| Male | 82 | IMF; BVVL | 5.269 | Male | 52 | Low value of RBC, HGB, HCT, MCV, MCH | 2.028 | Male | 44 | NA | 4.100 |
| Male | 56 | IMF; BVVL; PUS | 7.095 | Male | 27 | Low value of RBC, HGB, HCT, MCV, MCH | 2.544 | Male | 37 | NA | 3.100 |
| Male | 63 | IMF; BVVL; PUS | 8.844 | Male | 68 | Low value of RBC, HGB, HCT, MCV, MCH | 3.972 | Female | 49 | NA | 3.880 |
| Male | 58 | IMF; PUS | 5.599 | Male | 56 | Low value of RBC, HGB, HCT, MCV, MCH | 3.960 | Female | 56 | NA | 2.500 |
| Male | 78 | IMF; BVVL; PUS | 4.719 | Male | 70 | Low value of RBC, HGB, HCT, MCV, MCH | 6.396 | Female | 24 | NA | 2.390 |
| Female | 65 | IMF; BVVL | 5.709 | Female | 64 | Low value of RBC, HGB, HCT, MCV, MCH | 2.700 | Female | 33 | NA | 3.290 |
| Male | 55 | IMF; BVVL | 6.523 | Female | 71 | Low value of RBC, HGB, HCT, MCV, MCH | 2.628 | Male | 47 | NA | 3.490 |
| Male | 73 | IMF; PUS | 5.522 | Female | 67 | Low value of RBC, HGB, HCT, MCV, MCH | 2.208 | Male | 41 | NA | 2.380 |
| Female | 79 | BVVL; PUS | 5.346 | Female | 66 | Low value of RBC, HGB, HCT, MCV, MCH | 2.748 | Male | 24 | NA | 2.850 |
| Male | 78 | BVVL; PUS | 5.764 | Male | 60 | Low value of RBC, HGB, HCT, MCV, MCH | 3.708 | Male | 28 | NA | 2.500 |
| Male | 67 | BVVL; PUS | 6.512 | Male | 59 | Low value of RBC, HGB, HCT, MCV, MCH | 3.048 | Male | 29 | NA | 2.560 |
| Male | 67 | BVVL; PUS | 5.357 | Male | 54 | Low value of RBC, HGB, HCT, MCV, MCH | 5.280 | Male | 55 | NA | 2.740 |
| Female | 65 | IMF; BVVL; PUS | 5.093 | Female | 41 | Low value of RBC, HGB, HCT, MCV, MCH | 6.396 | Female | 51 | NA | 3.430 |
| Male | 54 | IMF; BVVL; PUS | 8.382 | Male | 46 | Low value of RBC, HGB, HCT, MCV, MCH | 4.404 | Female | 23 | NA | 2.740 |
| Male | 64 | IMF; BVVL; PUS | 6.941 | Male | 18 | Low value of RBC, HGB, HCT, MCV, MCH | 6.624 | Male | 57 | NA | 3.580 |
| Male | 61 | IMF; BVVL; PUS | 5.522 | Male | 38 | Low value of RBC, HGB, HCT, MCV, MCH | 2.700 | Female | 45 | NA | 4.930 |
| Female | 48 | BVVL; PUS | 5.005 | Male | 51 | Low value of RBC, HGB, HCT, MCV, MCH | 2.568 | Male | 29 | NA | 4.500 |
| Female | 51 | IMF; BVVL | 9.284 | Male | 23 | Low value of RBC, HGB, HCT, MCV, MCH | 3.552 | Male | 19 | NA | 3.300 |
| Male | 61 | BVVL; PUS | 5.907 | Male | 20 | Low value of RBC, HGB, HCT, MCV, MCH | 2.364 | Male | 58 | NA | 3.570 |
| Male | 67 | IMF; BVVL | 6.314 | Male | 26 | Low value of RBC, HGB, HCT, MCV, MCH | 3.360 | Female | 29 | NA | 3.100 |
| Female | 81 | IMF; BVVL; PUS | 6.028 | Male | 24 | Low value of RBC, HGB, HCT, MCV, MCH | 3.852 | Male | 25 | NA | 3.400 |
| Male | 45 | IMF; BVVL; PUS | 9.515 | Male | 23 | Low value of RBC, HGB, HCT, MCV, MCH | 3.012 | Female | 67 | NA | 3.000 |
| Male | 74 | IMF; BVVL; PUS | 8.448 | Male | 45 | Low value of RBC, HGB, HCT, MCV, MCH | 2.844 | Female | 46 | NA | 3.970 |
| Male | 79 | IMF; BVVL; PUS | 7.172 | Male | 19 | Low value of RBC, HGB, HCT, MCV, MCH | 2.388 | Female | 32 | NA | 2.630 |
| Male | 82 | IMF; BVVL | 7.458 | Male | 68 | Low value of RBC, HGB, HCT, MCV, MCH | 5.136 | Female | 65 | NA | 3.600 |
| Male | 62 | IMF; PUS | 9.625 | Male | 18 | Low value of RBC, HGB, HCT, MCV, MCH | 6.828 | Female | 27 | NA | 2.330 |
| Female | 80 | IMF; BVVL; PUS | 6.787 | Female | 16 | Low value of RBC, HGB, HCT, MCV, MCH | 5.136 | Female | 69 | NA | 2.070 |
| Female | 79 | IMF; BVVL; PUS | 6.743 | Female | 40 | Low value of RBC, HGB, HCT, MCV, MCH | 2.196 | Female | 19 | NA | 2.700 |
| Male | 69 | BVVL; PUS | 3.597 | Female | 39 | Low value of RBC, HGB, HCT, MCV, MCH | 5.292 | Male | 38 | NA | 3.200 |
| Male | 86 | IMF; BVVL; PUS | 3.377 | Male | 50 | Low value of RBC, HGB, HCT, MCV, MCH | 5.484 | Female | 32 | NA | 2.300 |
| Male | 42 | IMF; BVVL; PUS | 3.905 | Male | 72 | Low value of RBC, HGB, HCT, MCV, MCH | 2.760 | Male | 23 | NA | 2.390 |
| Male | 59 | BVVL; PUS | 5.984 | Male | 67 | Low value of RBC, HGB, HCT, MCV, MCH | 5.820 | Male | 26 | NA | 3.050 |
| Female | 79 | BVVL; PUS | 4.807 | Male | 60 | Low value of RBC, HGB, HCT, MCV, MCH | 2.916 | Male | 67 | NA | 2.740 |
| Female | 64 | BVVL; PUS | 3.014 | Male | 75 | Low value of RBC, HGB, HCT, MCV, MCH | 3.504 | Female | 43 | NA | 2.800 |
| Female | 63 | BVVL; PUS | 5.720 | Male | 25 | Low value of RBC, HGB, HCT, MCV, MCH | 2.088 | Male | 69 | NA | 2.380 |
| Female | 63 | IMF; BVVL | 9.515 | Female | 23 | Low value of RBC, HGB, HCT, MCV, MCH | 2.028 | Male | 75 | NA | 3.230 |
| Female | 39 | IMF; BVVL; PUS | 8.448 | Female | 76 | Low value of RBC, HGB, HCT, MCV, MCH | 8.808 | Male | 38 | NA | 2.480 |
| Male | 58 | IMF; BVVL; PUS | 7.172 | Male | 19 | Low value of RBC, HGB, HCT, MCV, MCH | 3.780 | Male | 73 | NA | 2.800 |
| Male | 70 | IMF; BVVL; PUS | 7.458 | Female | 22 | Low value of RBC, HGB, HCT, MCV, MCH | 3.624 | Male | 22 | NA | 2.900 |
| Male | 63 | IMF; BVVL; PUS | 9.625 | Female | 21 | Low value of RBC, HGB, HCT, MCV, MCH | 2.760 | Female | 37 | NA | 3.310 |
| Male | 41 | IMF; BVVL; PUS | 6.787 | Female | 20 | Low value of RBC, HGB, HCT, MCV, MCH | 5.988 | Female | 54 | NA | 2.480 |
| Female | 57 | IMF; BVVL | 6.743 | Female | 26 | Low value of RBC, HGB, HCT, MCV, MCH | 5.448 | Male | 30 | NA | 3.000 |
| Male | 28 | IMF; BVVL; PUS | 3.597 | Male | 19 | Low value of RBC, HGB, HCT, MCV, MCH | 5.436 | Female | 80 | NA | 2.490 |
| Female | 62 | BVVL; PUS | 3.377 | Female | 19 | Low value of RBC, HGB, HCT, MCV, MCH | 4.872 | Male | 33 | NA | 2.230 |
| Female | 72 | IMF; BVVL | 2.860 | Female | 21 | Low value of RBC, HGB, HCT, MCV, MCH | 3.156 | Female | 39 | NA | 2.700 |
| Male | 34 | IMF; BVVL; PUS | 3.278 | Male | 71 | Low value of RBC, HGB, HCT, MCV, MCH | 4.428 | Female | 30 | NA | 2.720 |
| Female | 70 | IMF; PUS | 3.916 | Male | 18 | Low value of RBC, HGB, HCT, MCV, MCH | 3.780 | Male | 78 | NA | 2.600 |
| Female | 78 | BVVL; PUS | 3.047 | Male | 20 | Low value of RBC, HGB, HCT, MCV, MCH | 3.036 | Male | 56 | NA | 2.610 |
| Male | 59 | IMF; PUS | 3.179 | Female | 22 | Low value of RBC, HGB, HCT, MCV, MCH | 3.972 | Female | 53 | NA | 2.930 |
| Male | 34 | IMF; BVVL; PUS | 5.412 | Female | 25 | Low value of RBC, HGB, HCT, MCV, MCH | 3.540 | Male | 34 | NA | 2.820 |
| Female | 75 | IMF; BVVL | 2.871 | Female | 69 | Low value of RBC, HGB, HCT, MCV, MCH | 1.824 | Female | 45 | NA | 2.410 |
| Male | 63 | BVVL; PUS | 4.532 | Female | 52 | Low value of RBC, HGB, HCT, MCV, MCH | 1.716 | Male | 70 | NA | 3.640 |
| Female | 77 | IMF; BVVL | 3.564 | Female | 72 | Low value of RBC, HGB, HCT, MCV, MCH | 9.864 | Female | 28 | NA | 2.740 |
| Female | 71 | IMF; BVVL; PUS | 4.950 | Female | 57 | Low value of RBC, HGB, HCT, MCV, MCH | 3.408 | Female | 51 | NA | 2.500 |
| Female | 78 | IMF; BVVL; PUS | 6.237 | Female | 56 | Low value of RBC, HGB, HCT, MCV, MCH | 4.656 | Female | 73 | NA | 2.800 |
| Male | 74 | IMF; BVVL; PUS | 5.159 | Male | 56 | Low value of RBC, HGB, HCT, MCV, MCH | 3.492 | Female | 27 | NA | 3.900 |
| Female | 73 | IMF; BVVL; PUS | 3.916 | Male | 32 | Low value of RBC, HGB, HCT, MCV, MCH | 4.968 | Female | 35 | NA | 2.580 |
| Male | 67 | IMF; BVVL; PUS | 4.719 | Male | 51 | Low value of RBC, HGB, HCT, MCV, MCH | 7.656 | Male | 56 | NA | 1.670 |
| Male | 66 | IMF; BVVL; PUS | 5.797 | Male | 63 | Low value of RBC, HGB, HCT, MCV, MCH | 11.304 | Male | 46 | NA | 3.100 |
| Female | 61 | IMF; PUS | 4.477 | Female | 56 | Low value of RBC, HGB, HCT, MCV, MCH | 3.420 | Female | 42 | NA | 3.000 |
| Male | 48 | BVVL; PUS | 6.589 | Male | 60 | Low value of RBC, HGB, HCT, MCV, MCH | 6.936 | Male | 38 | NA | 3.070 |
| Male | 53 | IMF; BVVL; PUS | 6.039 | Male | 74 | Low value of RBC, HGB, HCT, MCV, MCH | 4.968 | Male | 42 | NA | 2.740 |
| Female | 25 | IMF; BVVL; PUS | 4.059 | Male | 38 | Low value of RBC, HGB, HCT, MCV, MCH | 6.756 | Male | 48 | NA | 2.600 |
| Female | 45 | IMF; BVVL; PUS | 5.390 | Male | 67 | Low value of RBC, HGB, HCT, MCV, MCH | 8.784 | Female | 45 | NA | 3.310 |
| Male | 58 | IMF; BVVL | 3.729 | Male | 72 | Low value of RBC, HGB, HCT, MCV, MCH | 7.656 | Male | 25 | NA | 2.260 |
| Female | 30 | IMF; BVVL; PUS | 2.739 | Female | 75 | Low value of RBC, HGB, HCT, MCV, MCH | 3.372 | Male | 38 | NA | 2.900 |
| Male | 27 | BVVL; PUS | 2.816 | Male | 55 | Low value of RBC, HGB, HCT, MCV, MCH | 2.400 | Female | 21 | NA | 3.920 |
| Female | 33 | BVVL; PUS | 3.531 | Female | 73 | Low value of RBC, HGB, HCT, MCV, MCH | 7.068 | Female | 25 | NA | 2.580 |
| Female | 31 | BVVL; PUS | 5.665 | Female | 72 | Low value of RBC, HGB, HCT, MCV, MCH | 6.756 | Male | 27 | NA | 2.200 |
| Female | 30 | IMF; BVVL; PUS | 3.597 | Female | 62 | Low value of RBC, HGB, HCT, MCV, MCH | 10.884 | Female | 78 | NA | 2.390 |
| Female | 52 | IMF; BVVL; PUS | 3.773 | Female | 79 | Low value of RBC, HGB, HCT, MCV, MCH | 7.320 | Female | 20 | NA | 2.200 |
| Male | 26 | IMF; BVVL; PUS | 5.478 | Female | 35 | Low value of RBC, HGB, HCT, MCV, MCH | 6.936 | Female | 38 | NA | 2.300 |
| Male | 75 | IMF; BVVL; PUS | 4.565 | Female | 49 | Low value of RBC, HGB, HCT, MCV, MCH | 3.120 | Female | 47 | NA | 2.320 |
| Male | 25 | IMF; BVVL; PUS | 4.279 | Female | 56 | Low value of RBC, HGB, HCT, MCV, MCH | 2.184 | Female | 24 | NA | 2.380 |
| Male | 23 | IMF; BVVL; PUS | 4.169 | Female | 51 | Low value of RBC, HGB, HCT, MCV, MCH | 4.044 | Female | 40 | NA | 2.710 |
| Female | 47 | IMF; BVVL | 4.686 | Female | 71 | Low value of RBC, HGB, HCT, MCV, MCH | 2.184 | Male | 37 | NA | 2.500 |
| Female | 46 | IMF; BVVL; PUS | 3.201 | Female | 58 | Low value of RBC, HGB, HCT, MCV, MCH | 4.476 | Male | 68 | NA | 2.600 |
| Male | 57 | IMF; BVVL; PUS | 4.906 | Male | 48 | Low value of RBC, HGB, HCT, MCV, MCH | 2.880 | Male | 77 | NA | 2.670 |
| Female | 79 | BVVL; PUS | 5.456 | Male | 66 | Low value of RBC, HGB, HCT, MCV, MCH | 3.060 | Female | 22 | NA | 2.150 |
| Female | 74 | IMF; BVVL; PUS | 7.062 | Male | 72 | Low value of RBC, HGB, HCT, MCV, MCH | 2.760 | Female | 47 | NA | 3.370 |
| Female | 67 | IMF; BVVL; PUS | 6.270 | Female | 71 | Low value of RBC, HGB, HCT, MCV, MCH | 2.124 | Male | 35 | NA | 2.250 |
| Female | 82 | IMF; BVVL; PUS | 6.358 | Female | 60 | Low value of RBC, HGB, HCT, MCV, MCH | 2.256 | Female | 67 | NA | 3.080 |
| Male | 32 | IMF; BVVL; PUS | 7.348 | Male | 60 | Low value of RBC, HGB, HCT, MCV, MCH | 2.640 | Female | 77 | NA | 2.740 |
| Female | 30 | IMF; BVVL; PUS | 5.170 | Female | 58 | Low value of RBC, HGB, HCT, MCV, MCH | 2.952 | Male | 30 | NA | 2.540 |
| Male | 83 | IMF; BVVL; PUS | 5.456 | Male | 47 | Low value of RBC, HGB, HCT, MCV, MCH | 2.268 | Male | 61 | NA | 2.670 |
| Female | 26 | IMF; BVVL; PUS | 4.653 | Female | 57 | Low value of RBC, HGB, HCT, MCV, MCH | 3.432 | Male | 41 | NA | 2.540 |
| Male | 29 | IMF; BVVL; PUS | 4.587 | Female | 54 | Low value of RBC, HGB, HCT, MCV, MCH | 4.092 | Female | 35 | NA | 2.260 |
| Male | 28 | IMF; BVVL; PUS | 5.643 | Female | 41 | Low value of RBC, HGB, HCT, MCV, MCH | 1.884 | Male | 48 | NA | 3.390 |
| Male | 27 | IMF; BVVL; PUS | 5.940 | Male | 44 | Low value of RBC, HGB, HCT, MCV, MCH | 2.460 | Female | 26 | NA | 3.350 |
| Female | 33 | IMF; BVVL; PUS | 5.632 | Female | 54 | Low value of RBC, HGB, HCT, MCV, MCH | 1.440 | Female | 61 | NA | 2.700 |
| Male | 26 | IMF; BVVL; PUS | 4.092 | Male | 46 | Low value of RBC, HGB, HCT, MCV, MCH | 1.860 | Male | 23 | NA | 2.200 |
| Male | 26 | IMF; BVVL; PUS | 9.119 | Male | 64 | Low value of RBC, HGB, HCT, MCV, MCH | 2.040 | Female | 46 | NA | 2.610 |
| Male | 28 | IMF; BVVL; PUS | 7.370 | Male | 74 | Low value of RBC, HGB, HCT, MCV, MCH | 2.004 | Male | 80 | NA | 2.500 |
| Male | 78 | IMF; BVVL; PUS | 6.710 | Male | 77 | Low value of RBC, HGB, HCT, MCV, MCH | 9.636 | Male | 65 | NA | 2.490 |
| Female | 25 | IMF; BVVL; PUS | 5.665 | Female | 66 | Low value of RBC, HGB, HCT, MCV, MCH | 2.784 | Female | 42 | NA | 2.850 |
| Male | 27 | IMF; BVVL; PUS | 8.151 | Male | 63 | Low value of RBC, HGB, HCT, MCV, MCH | 5.064 | Female | 74 | NA | 2.410 |
| Female | 29 | IMF; BVVL; PUS | 5.104 | Male | 47 | Low value of RBC, HGB, HCT, MCV, MCH | 2.220 | Female | 58 | NA | 2.850 |
| Female | 32 | IMF; BVVL; PUS | 5.214 | Male | 47 | Low value of RBC, HGB, HCT, MCV, MCH | 3.408 | Female | 37 | NA | 2.430 |
| Male | 76 | IMF; BVVL; PUS | 6.347 | Male | 80 | Low value of RBC, HGB, HCT, MCV, MCH | 2.772 | Female | 25 | NA | 3.830 |
| Female | 70 | IMF; BVVL; PUS | 5.236 | Male | 59 | Low value of RBC, HGB, HCT, MCV, MCH | 2.964 | Male | 63 | NA | 2.530 |
| Female | 27 | IMF; BVVL; PUS | 4.092 | Male | 49 | Low value of RBC, HGB, HCT, MCV, MCH | 5.244 | Male | 24 | NA | 2.960 |
| Male | 69 | IMF; BVVL; PUS | 6.083 | Male | 51 | Low value of RBC, HGB, HCT, MCV, MCH | 2.820 | Male | 44 | NA | 2.540 |
| Female | 68 | IMF; BVVL; PUS | 5.819 | Female | 61 | Low value of RBC, HGB, HCT, MCV, MCH | 2.880 | Female | 26 | NA | 2.620 |
| Female | 37 | IMF; BVVL; PUS | 3.652 | Female | 59 | Low value of RBC, HGB, HCT, MCV, MCH | 3.348 | Male | 35 | NA | 3.080 |
| Male | 63 | IMF; BVVL; PUS | 7.931 | Female | 80 | Low value of RBC, HGB, HCT, MCV, MCH | 8.832 | Female | 45 | NA | 2.970 |
| Male | 77 | IMF; BVVL; PUS | 5.698 | Female | 39 | Low value of RBC, HGB, HCT, MCV, MCH | 6.120 | Female | 23 | NA | 2.500 |
| Female | 68 | IMF; BVVL; PUS | 6.116 | Female | 80 | Low value of RBC, HGB, HCT, MCV, MCH | 7.044 | Female | 23 | NA | 2.600 |
| Male | 66 | IMF; BVVL; PUS | 5.357 | Male | 74 | Low value of RBC, HGB, HCT, MCV, MCH | 2.844 | Male | 25 | NA | 3.000 |
| Female | 71 | IMF; BVVL; PUS | 5.104 | Male | 81 | Low value of RBC, HGB, HCT, MCV, MCH | 5.124 | Male | 43 | NA | 3.760 |
| Male | 71 | IMF; BVVL; PUS | 4.631 | Female | 64 | Low value of RBC, HGB, HCT, MCV, MCH | 2.172 | Female | 23 | NA | 3.260 |
| Female | 64 | IMF; BVVL; PUS | 7.997 | Female | 69 | Low value of RBC, HGB, HCT, MCV, MCH | 3.192 | Male | 34 | NA | 2.100 |
| Female | 65 | IMF; BVVL; PUS | 4.422 | Female | 69 | Low value of RBC, HGB, HCT, MCV, MCH | 2.964 | Male | 71 | NA | 3.400 |
| Female | 61 | IMF; BVVL; PUS | 5.929 | Female | 62 | Low value of RBC, HGB, HCT, MCV, MCH | 8.328 | Male | 49 | NA | 2.400 |
| Male | 51 | IMF; BVVL; PUS | 3.839 | Female | 63 | Low value of RBC, HGB, HCT, MCV, MCH | 4.968 | Female | 50 | NA | 2.100 |
| Female | 53 | IMF; BVVL; PUS | 5.786 | Female | 74 | Low value of RBC, HGB, HCT, MCV, MCH | 4.500 | Female | 31 | NA | 3.470 |
| Male | 63 | IMF; BVVL; PUS | 3.267 | Male | 62 | Low value of RBC, HGB, HCT, MCV, MCH | 2.544 | Male | 59 | NA | 3.720 |
| Male | 61 | IMF; BVVL; PUS | 7.216 | Male | 73 | Low value of RBC, HGB, HCT, MCV, MCH | 12.108 | Male | 18 | NA | 2.330 |
| Male | 82 | IMF; BVVL; PUS | 4.268 | Male | 68 | Low value of RBC, HGB, HCT, MCV, MCH | 2.196 | Female | 39 | NA | 1.900 |
| Male | 41 | IMF; BVVL; PUS | 6.358 | Male | 25 | Low value of RBC, HGB, HCT, MCV, MCH | 2.652 | Male | 25 | NA | 2.600 |
| Male | 82 | IMF; BVVL; PUS | 2.563 | Male | 67 | Low value of RBC, HGB, HCT, MCV, MCH | 3.624 | Female | 48 | NA | 2.810 |
| Male | 76 | IMF; BVVL; PUS | 4.763 | Male | 66 | Low value of RBC, HGB, HCT, MCV, MCH | 4.464 | Female | 50 | NA | 3.390 |
| Female | 83 | IMF; BVVL; PUS | 3.696 | Male | 35 | Low value of RBC, HGB, HCT, MCV, MCH | 4.440 | Male | 53 | NA | 2.520 |
| Male | 76 | IMF; BVVL; PUS | 3.938 | Male | 61 | Low value of RBC, HGB, HCT, MCV, MCH | 4.596 | Female | 24 | NA | 2.540 |
| Male | 64 | IMF; BVVL; PUS | 2.662 | Male | 75 | Low value of RBC, HGB, HCT, MCV, MCH | 2.544 | Male | 28 | NA | 2.180 |
| Male | 75 | IMF; BVVL; PUS | 3.146 | Male | 66 | Low value of RBC, HGB, HCT, MCV, MCH | 4.440 | Female | 36 | NA | 2.390 |
| Female | 73 | IMF; BVVL; PUS | 3.410 | Male | 71 | Low value of RBC, HGB, HCT, MCV, MCH | 2.616 | Female | 37 | NA | 2.770 |
| Female | 47 | IMF; BVVL; PUS | 4.653 | Male | 57 | Low value of RBC, HGB, HCT, MCV, MCH | 3.912 | Male | 28 | NA | 2.360 |
| Male | 69 | IMF; BVVL; PUS | 4.994 | Male | 30 | Low value of RBC, HGB, HCT, MCV, MCH | 4.692 | Female | 50 | NA | 2.480 |
| Male | 63 | IMF; BVVL; PUS | 4.576 | Male | 62 | Low value of RBC, HGB, HCT, MCV, MCH | 4.188 | Male | 24 | NA | 3.920 |
| Male | 75 | IMF; BVVL; PUS | 3.685 | Male | 48 | Low value of RBC, HGB, HCT, MCV, MCH | 2.328 | Female | 53 | NA | 1.800 |
| Female | 48 | IMF; BVVL; PUS | 5.181 | Female | 23 | Low value of RBC, HGB, HCT, MCV, MCH | 3.252 | Female | 25 | NA | 3.020 |
| Female | 60 | IMF; BVVL; PUS | 3.751 | Female | 44 | Low value of RBC, HGB, HCT, MCV, MCH | 2.904 | Female | 24 | NA | 2.410 |
| Male | 37 | IMF; BVVL; PUS | 2.387 | Male | 63 | Low value of RBC, HGB, HCT, MCV, MCH | 9.012 | Female | 42 | NA | 5.000 |
| Female | 64 | IMF; BVVL; PUS | 3.718 | Female | 41 | Low value of RBC, HGB, HCT, MCV, MCH | 3.696 | Male | 49 | NA | 2.700 |
| Male | 29 | IMF; BVVL; PUS | 5.874 | Female | 70 | Low value of RBC, HGB, HCT, MCV, MCH | 2.724 | Female | 55 | NA | 4.940 |
| Male | 59 | IMF; BVVL; PUS | 2.827 | Female | 35 | Low value of RBC, HGB, HCT, MCV, MCH | 3.360 | Male | 34 | NA | 3.110 |
| Female | 68 | IMF; BVVL; PUS | 3.861 | Female | 62 | Low value of RBC, HGB, HCT, MCV, MCH | 9.300 | Female | 18 | NA | 2.890 |
| Female | 59 | IMF; BVVL; PUS | 3.300 | Female | 27 | Low value of RBC, HGB, HCT, MCV, MCH | 2.268 | Female | 64 | NA | 2.850 |
| Female | 32 | IMF; BVVL; PUS | 3.058 | Female | 57 | Low value of RBC, HGB, HCT, MCV, MCH | 4.116 | Male | 63 | NA | 3.230 |
| Male | 64 | IMF; BVVL; PUS | 5.544 | Male | 66 | Low value of RBC, HGB, HCT, MCV, MCH | 2.592 | Female | 29 | NA | 2.880 |
| Female | 50 | IMF; BVVL; PUS | 2.805 | Male | 45 | Low value of RBC, HGB, HCT, MCV, MCH | 5.304 | Female | 45 | NA | 1.960 |
| Male | 25 | IMF; BVVL; PUS | 2.849 | Male | 67 | Low value of RBC, HGB, HCT, MCV, MCH | 2.028 | Female | 35 | NA | 2.070 |
| Female | 46 | IMF; BVVL; PUS | 4.158 | Female | 61 | Low value of RBC, HGB, HCT, MCV, MCH | 2.844 | Male | 68 | NA | 2.290 |
| Male | 65 | IMF; BVVL; PUS | 3.234 | Male | 73 | Low value of RBC, HGB, HCT, MCV, MCH | 2.388 | Male | 37 | NA | 2.180 |
| Male | 43 | IMF; BVVL; PUS | 5.478 | Female | 46 | Low value of RBC, HGB, HCT, MCV, MCH | 2.640 | Male | 59 | NA | 2.070 |
| Female | 72 | IMF; BVVL; PUS | 4.466 | Female | 58 | Low value of RBC, HGB, HCT, MCV, MCH | 2.340 | Male | 61 | NA | 2.100 |
| Female | 30 | IMF; BVVL; PUS | 3.795 | Female | 28 | Low value of RBC, HGB, HCT, MCV, MCH | 4.920 | Male | 19 | NA | 2.290 |
| Female | 42 | IMF; BVVL; PUS | 5.709 | Female | 51 | Low value of RBC, HGB, HCT, MCV, MCH | 1.992 | Female | 51 | NA | 2.270 |
| Male | 35 | IMF; BVVL; PUS | 3.113 | Male | 28 | Low value of RBC, HGB, HCT, MCV, MCH | 9.864 | Female | 23 | NA | 3.370 |
| Female | 42 | IMF; BVVL; PUS | 3.575 | Female | 36 | Low value of RBC, HGB, HCT, MCV, MCH | 8.724 | Male | 30 | NA | 2.250 |
| Male | 40 | IMF; BVVL; PUS | 5.170 | Female | 31 | Low value of RBC, HGB, HCT, MCV, MCH | 3.792 | Male | 67 | NA | 2.180 |
| Male | 28 | IMF; BVVL; PUS | 3.278 | Male | 54 | Low value of RBC, HGB, HCT, MCV, MCH | 1.824 | Female | 51 | NA | 2.300 |
| Female | 40 | IMF; BVVL; PUS | 2.530 | Male | 45 | Low value of RBC, HGB, HCT, MCV, MCH | 2.028 | Male | 22 | NA | 2.380 |
| Female | 25 | IMF; BVVL; PUS | 3.575 | Female | 76 | Low value of RBC, HGB, HCT, MCV, MCH | 2.544 | Female | 70 | NA | 2.070 |
| Female | 40 | IMF; BVVL; PUS | 4.378 | Female | 54 | Low value of RBC, HGB, HCT, MCV, MCH | 3.972 | Female | 38 | NA | 2.380 |
| Male | 43 | IMF; BVVL; PUS | 3.597 | Male | 76 | Low value of RBC, HGB, HCT, MCV, MCH | 2.124 | Female | 40 | NA | 2.210 |
| Male | 41 | IMF; BVVL; PUS | 3.905 | Female | 40 | Low value of RBC, HGB, HCT, MCV, MCH | 3.960 | Female | 53 | NA | 2.300 |
| Male | 68 | IMF; BVVL; PUS | 2.816 | Female | 52 | Low value of RBC, HGB, HCT, MCV, MCH | 6.396 | Female | 35 | NA | 3.350 |
| Male | 57 | IMF; BVVL; PUS | 3.795 | Male | 24 | Low value of RBC, HGB, HCT, MCV, MCH | 2.700 | Female | 51 | NA | 5.400 |
| Male | 80 | IMF; BVVL; PUS | 3.124 | Male | 40 | Low value of RBC, HGB, HCT, MCV, MCH | 2.628 | Female | 50 | NA | 1.880 |
| Female | 37 | IMF; BVVL; PUS | 2.959 | Female | 33 | Low value of RBC, HGB, HCT, MCV, MCH | 2.208 | Female | 28 | NA | 2.380 |
| Male | 45 | IMF; BVVL; PUS | 4.631 | Male | 40 | Low value of RBC, HGB, HCT, MCV, MCH | 2.748 | Female | 37 | NA | 2.390 |
| Male | 67 | IMF; BVVL; PUS | 6.567 | Male | 38 | Low value of RBC, HGB, HCT, MCV, MCH | 3.708 | Female | 41 | NA | 2.460 |
| Female | 53 | IMF; BVVL; PUS | 4.037 | Male | 26 | Low value of RBC, HGB, HCT, MCV, MCH | 3.048 | Female | 56 | NA | 2.490 |
| Female | 30 | IMF; BVVL; PUS | 2.750 | Male | 38 | Low value of RBC, HGB, HCT, MCV, MCH | 2.160 | Female | 37 | NA | 2.410 |
| Male | 38 | IMF; BVVL; PUS | 6.116 | Male | 66 | Low value of RBC, HGB, HCT, MCV, MCH | 2.100 | Male | 38 | NA | 2.500 |
| Male | 33 | IMF; BVVL; PUS | 3.410 | Male | 55 | Low value of RBC, HGB, HCT, MCV, MCH | 5.136 | Male | 26 | NA | 2.360 |
| Male | 56 | IMF; BVVL; PUS | 4.125 | Male | 78 | Low value of RBC, HGB, HCT, MCV, MCH | 6.828 | Female | 25 | NA | 3.830 |
| Male | 47 | IMF; BVVL; PUS | 2.750 | Female | 35 | Low value of RBC, HGB, HCT, MCV, MCH | 2.748 | Female | 38 | NA | 2.240 |
| Male | 78 | IMF; BVVL; PUS | 3.025 | Female | 43 | Low value of RBC, HGB, HCT, MCV, MCH | 5.280 | Female | 48 | NA | 2.740 |
| Female | 56 | IMF; BVVL; PUS | 3.091 | Female | 65 | Low value of RBC, HGB, HCT, MCV, MCH | 6.396 | Female | 39 | NA | 1.490 |
| Male | 78 | IMF; BVVL; PUS | 3.245 | Male | 23 | Low value of RBC, HGB, HCT, MCV, MCH | 4.404 | Female | 28 | NA | 2.620 |
| Male | 42 | IMF; BVVL; PUS | 3.740 | Male | 38 | Low value of RBC, HGB, HCT, MCV, MCH | 6.624 | Female | 47 | NA | 3.080 |
| Female | 54 | IMF; BVVL; PUS | 4.829 | Male | 41 | Low value of RBC, HGB, HCT, MCV, MCH | 2.700 | Female | 29 | NA | 2.740 |
| Female | 26 | IMF; BVVL; PUS | 3.553 | Male | 39 | Low value of RBC, HGB, HCT, MCV, MCH | 2.220 | Female | 43 | NA | 2.380 |
| Male | 75 | IMF; BVVL; PUS | 4.796 | Male | 73 | Low value of RBC, HGB, HCT, MCV, MCH | 5.136 | Male | 67 | NA | 2.290 |
| Male | 45 | IMF; BVVL; PUS | 4.378 | Male | 43 | Low value of RBC, HGB, HCT, MCV, MCH | 2.196 | Male | 50 | NA | 2.220 |
| Male | 38 | IMF; BVVL; PUS | 3.718 | Male | 36 | Low value of RBC, HGB, HCT, MCV, MCH | 2.136 | Female | 28 | NA | 2.020 |
| Male | 47 | IMF; BVVL; PUS | 7.557 | Male | 45 | Low value of RBC, HGB, HCT, MCV, MCH | 2.568 | Male | 56 | NA | 3.260 |
| Male | 52 | IMF; BVVL; PUS | 2.387 | Male | 50 | Low value of RBC, HGB, HCT, MCV, MCH | 3.552 | Female | 41 | NA | 2.230 |
| Female | 34 | IMF; BVVL; PUS | 2.695 | Male | 32 | Low value of RBC, HGB, HCT, MCV, MCH | 2.364 | Female | 58 | NA | 2.280 |
| Male | 39 | IMF; BVVL; PUS | 4.719 | Male | 37 | Low value of RBC, HGB, HCT, MCV, MCH | 3.360 | Male | 49 | NA | 2.270 |
| Male | 36 | IMF; BVVL; PUS | 3.212 | Male | 34 | Low value of RBC, HGB, HCT, MCV, MCH | 3.852 | Female | 27 | NA | 2.090 |
| Female | 22 | IMF; BVVL; PUS | 2.750 | Female | 20 | Low value of RBC, HGB, HCT, MCV, MCH | 3.012 | Female | 20 | NA | 3.470 |
| Male | 35 | IMF; BVVL; PUS | 2.244 | Female | 33 | Low value of RBC, HGB, HCT, MCV, MCH | 1.788 | Male | 32 | NA | 3.720 |
| Female | 50 | IMF; BVVL; PUS | 3.179 | Female | 48 | Low value of RBC, HGB, HCT, MCV, MCH | 1.956 | Male | 45 | NA | 2.330 |
| Female | 31 | IMF; BVVL; PUS | 3.707 | Female | 29 | Low value of RBC, HGB, HCT, MCV, MCH | 5.292 | Female | 67 | NA | 2.070 |
| Male | 46 | IMF; BVVL; PUS | 2.992 | Male | 44 | Low value of RBC, HGB, HCT, MCV, MCH | 5.484 | Female | 60 | NA | 2.600 |
| Male | 38 | IMF; BVVL; PUS | 3.553 | Male | 36 | Low value of RBC, HGB, HCT, MCV, MCH | 2.760 | Female | 55 | NA | 2.810 |
| Female | 42 | IMF; BVVL; PUS | 4.213 | Female | 40 | Low value of RBC, HGB, HCT, MCV, MCH | 5.820 | Male | 24 | NA | 3.390 |
| Male | 34 | IMF; BVVL; PUS | 3.641 | Male | 32 | Low value of RBC, HGB, HCT, MCV, MCH | 1.992 | Female | 21 | NA | 2.520 |
| Male | 48 | IMF; BVVL; PUS | 2.530 | Male | 46 | Low value of RBC, HGB, HCT, MCV, MCH | 2.916 | Male | 22 | NA | 2.540 |
| Female | 32 | IMF; BVVL; PUS | 6.061 | Female | 30 | Low value of RBC, HGB, HCT, MCV, MCH | 1.764 | Female | 22 | NA | 2.420 |
| Male | 79 | IMF; BVVL; PUS | 2.981 | Male | 77 | Low value of RBC, HGB, HCT, MCV, MCH | 1.824 | Male | 39 | NA | 2.240 |
| Female | 19 | IMF; BVVL; PUS | 3.509 | Female | 17 | Low value of RBC, HGB, HCT, MCV, MCH | 3.504 | Female | 31 | NA | 2.770 |
| Male | 49 | IMF; BVVL; PUS | 2.684 | Male | 47 | Low value of RBC, HGB, HCT, MCV, MCH | 1.536 | Male | 48 | NA | 2.360 |
| Male | 28 | IMF; BVVL; PUS | 3.564 | Female | 26 | Low value of RBC, HGB, HCT, MCV, MCH | 2.088 | Female | 70 | NA | 2.480 |
| Male | 30 | IMF; BVVL; PUS | 5.137 | Female | 28 | Low value of RBC, HGB, HCT, MCV, MCH | 2.028 | Female | 78 | NA | 3.920 |
| Male | 67 | IMF; BVVL; PUS | 3.564 | Male | 65 | Low value of RBC, HGB, HCT, MCV, MCH | 8.808 | Male | 31 | NA | 1.960 |
| Female | 45 | IMF; BVVL; PUS | 3.201 | Male | 43 | Low value of RBC, HGB, HCT, MCV, MCH | 3.780 | Male | 48 | NA | 3.020 |
| Female | 18 | IMF; BVVL; PUS | 3.256 | Male | 16 | Low value of RBC, HGB, HCT, MCV, MCH | 3.624 | Male | 67 | NA | 2.410 |
| Male | 67 | IMF; BVVL; PUS | 3.773 | Male | 65 | Low value of RBC, HGB, HCT, MCV, MCH | 2.760 | Female | 22 | NA | 2.290 |
| Female | 40 | IMF; BVVL; PUS | 4.323 | Female | 38 | Low value of RBC, HGB, HCT, MCV, MCH | 5.988 | Male | 44 | NA | 2.130 |
| Male | 79 | IMF; BVVL; PUS | 4.202 | Male | 77 | Low value of RBC, HGB, HCT, MCV, MCH | 1.608 | Female | 49 | NA | 4.940 |
| Male | 62 | IMF; BVVL; PUS | 4.873 | Male | 60 | Low value of RBC, HGB, HCT, MCV, MCH | 2.088 | Male | 35 | NA | 3.110 |
| Male | 35 | IMF; BVVL; PUS | 4.169 | Male | 33 | Low value of RBC, HGB, HCT, MCV, MCH | 1.440 | Male | 27 | NA | 2.890 |
| Female | 40 | IMF; BVVL; PUS | 7.040 | Female | 38 | Low value of RBC, HGB, HCT, MCV, MCH | 5.448 | Male | 38 | NA | 2.850 |
| Male | 44 | IMF; BVVL; PUS | 3.465 | Female | 42 | Low value of RBC, HGB, HCT, MCV, MCH | 1.248 | Female | 23 | NA | 3.230 |
| Male | 49 | IMF; BVVL; PUS | 6.281 | Male | 47 | Low value of RBC, HGB, HCT, MCV, MCH | 2.112 | Male | 60 | NA | 2.880 |
| Male | 83 | IMF; BVVL; PUS | 4.257 | Female | 81 | Low value of RBC, HGB, HCT, MCV, MCH | 3.156 | Female | 70 | NA | 2.140 |
| Male | 47 | IMF; BVVL; PUS | 6.248 | Female | 45 | Low value of RBC, HGB, HCT, MCV, MCH | 1.896 | Female | 27 | NA | 2.110 |
| Male | 48 | IMF; BVVL; PUS | 3.652 | Female | 46 | Low value of RBC, HGB, HCT, MCV, MCH | 5.436 | Female | 24 | NA | 1.730 |
| Male | 79 | IMF; BVVL; PUS | 4.092 | Female | 77 | Low value of RBC, HGB, HCT, MCV, MCH | 4.872 | Female | 39 | NA | 2.410 |
| Female | 93 | IMF; BVVL | 5.093 | Female | 91 | Low value of RBC, HGB, HCT, MCV, MCH | 1.920 | Female | 27 | NA | 2.423 |
| Male | 59 | IMF; BVVL; PUS | 4.290 | Male | 57 | Low value of RBC, HGB, HCT, MCV, MCH | 4.428 | Male | 45 | NA | 2.790 |
| Female | 84 | IMF; BVVL; PUS | 3.509 | Female | 82 | Low value of RBC, HGB, HCT, MCV, MCH | 3.780 | Male | 54 | NA | 2.240 |
| Male | 59 | IMF; BVVL; PUS | 3.047 | Male | 57 | Low value of RBC, HGB, HCT, MCV, MCH | 3.036 | Female | 77 | NA | 2.610 |
| Female | 74 | IMF; BVVL; PUS | 3.938 | Female | 72 | Low value of RBC, HGB, HCT, MCV, MCH | 3.972 | Female | 84 | NA | 2.489 |
| Male | 64 | IMF; BVVL; PUS | 2.695 | Male | 62 | Low value of RBC, HGB, HCT, MCV, MCH | 3.540 | Female | 21 | NA | 2.790 |
| Male | 47 | IMF; BVVL; PUS | 4.818 | Male | 45 | Low value of RBC, HGB, HCT, MCV, MCH | 1.932 | Female | 62 | NA | 2.256 |
| Male | 48 | IMF; BVVL; PUS | 5.104 | Male | 46 | Low value of RBC, HGB, HCT, MCV, MCH | 2.064 | Female | 52 | NA | 2.610 |
| Female | 76 | IMF; BVVL; PUS | 4.796 | Female | 74 | Low value of RBC, HGB, HCT, MCV, MCH | 1.524 | Male | 23 | NA | 2.489 |
| Male | 62 | IMF; BVVL; PUS | 2.211 | Male | 60 | Low value of RBC, HGB, HCT, MCV, MCH | 1.824 | Female | 68 | NA | 3.350 |
| Female | 74 | IMF; BVVL; PUS | 2.277 | Female | 72 | Low value of RBC, HGB, HCT, MCV, MCH | 1.716 | Female | 38 | NA | 5.400 |
| Female | 59 | IMF; BVVL; PUS | 4.829 | Female | 57 | Low value of RBC, HGB, HCT, MCV, MCH | 1.356 | Female | 50 | NA | 1.730 |
| Female | 19 | IMF; BVVL; PUS | 4.136 | Female | 17 | Low value of RBC, HGB, HCT, MCV, MCH | 9.864 | Male | 47 | NA | 2.410 |
| Male | 80 | IMF; BVVL; PUS | 3.080 | Male | 78 | Low value of RBC, HGB, HCT, MCV, MCH | 3.408 | Female | 50 | NA | 6.000 |
| Male | 48 | IMF; BVVL; PUS | 3.784 | Female | 46 | Low value of RBC, HGB, HCT, MCV, MCH | 4.656 | Male | 47 | NA | 1.490 |
| Male | 66 | IMF; BVVL; PUS | 3.861 | Female | 64 | Low value of RBC, HGB, HCT, MCV, MCH | 3.492 | Female | 38 | NA | 2.070 |
| Female | 93 | IMF; BVVL; PUS | 3.366 | Female | 91 | Low value of RBC, HGB, HCT, MCV, MCH | 4.968 | Female | 55 | NA | 2.410 |
| Female | 46 | IMF; BVVL; PUS | 2.970 | Female | 44 | Low value of RBC, HGB, HCT, MCV, MCH | 7.656 | Female | 47 | NA | 2.180 |
| Male | 70 | IMF; BVVL; PUS | 4.279 | Male | 68 | Low value of RBC, HGB, HCT, MCV, MCH | 11.304 | Male | 71 | NA | 2.230 |
| Male | 46 | IMF; BVVL; PUS | 3.916 | Female | 44 | Low value of RBC, HGB, HCT, MCV, MCH | 3.420 | Female | 35 | NA | 3.830 |
| Male | 33 | IMF; BVVL; PUS | 3.674 | Female | 31 | Low value of RBC, HGB, HCT, MCV, MCH | 6.936 | Female | 33 | NA | 2.640 |
| Female | 63 | IMF; BVVL; PUS | 2.695 | Female | 61 | Low value of RBC, HGB, HCT, MCV, MCH | 4.968 | Female | 67 | NA | 2.410 |
| Male | 55 | IMF; BVVL; PUS | 3.531 | Male | 53 | Low value of RBC, HGB, HCT, MCV, MCH | 6.756 | Female | 40 | NA | 2.830 |
| Male | 80 | IMF; BVVL; PUS | 5.027 | Male | 78 | Low value of RBC, HGB, HCT, MCV, MCH | 8.784 | Female | 78 | NA | 3.760 |
| Female | 65 | IMF; BVVL; PUS | 3.652 | Female | 63 | Low value of RBC, HGB, HCT, MCV, MCH | 7.656 | Female | 36 | NA | 3.260 |
| Female | 84 | IMF; BVVL; PUS | 4.015 | Female | 82 | Low value of RBC, HGB, HCT, MCV, MCH | 3.372 | Female | 43 | NA | 2.100 |
| Female | 67 | IMF; BVVL; PUS | 3.146 | Female | 65 | Low value of RBC, HGB, HCT, MCV, MCH | 2.400 | Female | 21 | NA | 3.960 |
| Male | 55 | IMF; BVVL; PUS | 5.093 | Male | 53 | Low value of RBC, HGB, HCT, MCV, MCH | 7.068 | Female | 50 | NA | 3.260 |
| Male | 65 | IMF; BVVL; PUS | 3.355 | Female | 63 | Low value of RBC, HGB, HCT, MCV, MCH | 6.756 | Female | 37 | NA | 2.100 |
| Female | 50 | IMF; BVVL; PUS | 4.488 | Female | 48 | Low value of RBC, HGB, HCT, MCV, MCH | 10.884 | Female | 29 | NA | 3.760 |
| Male | 73 | IMF; BVVL; PUS | 3.861 | Female | 71 | Low value of RBC, HGB, HCT, MCV, MCH | 7.320 | Female | 52 | NA | 3.260 |
| Female | 80 | IMF; BVVL; PUS | 4.125 | Female | 78 | Low value of RBC, HGB, HCT, MCV, MCH | 6.936 | Male | 41 | NA | 2.400 |
| Male | 75 | IMF; BVVL; PUS | 3.839 | Male | 73 | Low value of RBC, HGB, HCT, MCV, MCH | 5.244 | Male | 28 | NA | 2.440 |
| Male | 76 | IMF; BVVL; PUS | 3.938 | Female | 74 | Low value of RBC, HGB, HCT, MCV, MCH | 2.820 | Female | 30 | NA | 2.740 |
| Male | 74 | IMF; BVVL; PUS | 2.508 | Female | 72 | Low value of RBC, HGB, HCT, MCV, MCH | 2.880 | Male | 25 | NA | 2.290 |
| Male | 80 | IMF; BVVL; PUS | 4.356 | Female | 78 | Low value of RBC, HGB, HCT, MCV, MCH | 3.348 | Female | 41 | NA | 3.470 |
| Male | 30 | IMF; BVVL; PUS | 3.850 | Female | 28 | Low value of RBC, HGB, HCT, MCV, MCH | 8.832 | Male | 42 | NA | 3.720 |
| Male | 61 | IMF; BVVL; PUS | 9.801 | Male | 59 | Low value of RBC, HGB, HCT, MCV, MCH | 6.120 | Male | 5 | NA | 2.330 |
| Male | 70 | IMF; BVVL; PUS | 3.300 | Male | 68 | Low value of RBC, HGB, HCT, MCV, MCH | 7.044 | Female | 31 | NA | 2.100 |
| Female | 75 | IMF; BVVL; PUS | 2.871 | Female | 73 | Low value of RBC, HGB, HCT, MCV, MCH | 2.844 | Female | 42 | NA | 2.600 |
| Male | 62 | IMF; BVVL; PUS | 3.421 | Male | 60 | Low value of RBC, HGB, HCT, MCV, MCH | 5.124 | Female | 40 | NA | 2.810 |
| Female | 74 | IMF; BVVL; PUS | 5.302 | Female | 72 | Low value of RBC, HGB, HCT, MCV, MCH | 3.192 | Male | 26 | NA | 3.390 |
| Male | 50 | IMF; BVVL; PUS | 2.904 | Male | 48 | Low value of RBC, HGB, HCT, MCV, MCH | 2.964 | Male | 56 | NA | 2.520 |
| Male | 86 | IMF; BVVL; PUS | 4.532 | Male | 84 | Low value of RBC, HGB, HCT, MCV, MCH | 8.328 | Male | 33 | NA | 2.540 |
| Male | 54 | IMF; BVVL; PUS | 4.125 | Male | 52 | Low value of RBC, HGB, HCT, MCV, MCH | 4.968 | Female | 6 | NA | 3.040 |
| Female | 58 | IMF; BVVL; PUS | 2.750 | Male | 56 | Low value of RBC, HGB, HCT, MCV, MCH | 4.500 | Female | 47 | NA | 2.850 |
| Female | 67 | IMF; BVVL; PUS | 3.025 | Male | 65 | Low value of RBC, HGB, HCT, MCV, MCH | 2.196 | Female | 45 | NA | 2.770 |
| Male | 77 | IMF; BVVL; PUS | 3.091 | Female | 75 | Low value of RBC, HGB, HCT, MCV, MCH | 2.652 | Male | 4 | NA | 3.290 |
| Male | 71 | IMF; BVVL; PUS | 3.245 | Female | 69 | Low value of RBC, HGB, HCT, MCV, MCH | 3.624 | Female | 3 | NA | 2.480 |
| Female | 77 | IMF; BVVL; PUS | 3.740 | Male | 75 | Low value of RBC, HGB, HCT, MCV, MCH | 4.464 | Female | 50 | NA | 3.920 |
| Female | 79 | IMF; BVVL; PUS | 2.453 | Male | 77 | Low value of RBC, HGB, HCT, MCV, MCH | 4.440 | Female | 80 | NA | 2.300 |
| Male | 88 | IMF; BVVL; PUS | 2.552 | Male | 86 | Low value of RBC, HGB, HCT, MCV, MCH | 4.596 | Male | 56 | NA | 3.020 |
| Male | 54 | IMF; BVVL; PUS | 4.796 | Male | 52 | Low value of RBC, HGB, HCT, MCV, MCH | 4.440 | Female | 53 | NA | 2.200 |
| Female | 67 | IMF; BVVL; PUS | 4.378 | Female | 65 | Low value of RBC, HGB, HCT, MCV, MCH | 2.616 | Male | 45 | NA | 3.300 |
| Male | 66 | IMF; BVVL; PUS | 3.718 | Female | 64 | Low value of RBC, HGB, HCT, MCV, MCH | 3.912 | Male | 39 | NA | 1.900 |
| Male | 34 | IMF; BVVL; PUS | 7.557 | Female | 32 | Low value of RBC, HGB, HCT, MCV, MCH | 4.692 | Female | 32 | NA | 4.940 |
| Male | 69 | IMF; BVVL; PUS | 2.387 | Female | 67 | Low value of RBC, HGB, HCT, MCV, MCH | 4.188 | Female | 31 | NA | 3.110 |
| Male | 28 | IMF; BVVL; PUS | 2.200 | Female | 26 | Low value of RBC, HGB, HCT, MCV, MCH | 2.328 | Female | 26 | NA | 2.890 |
| Male | 78 | IMF; BVVL; PUS | 4.719 | Male | 76 | Low value of RBC, HGB, HCT, MCV, MCH | 3.252 | Female | 42 | NA | 2.850 |
| Male | 78 | IMF; BVVL; PUS | 3.212 | Male | 76 | Low value of RBC, HGB, HCT, MCV, MCH | 2.904 | Female | 28 | NA | 3.230 |
| Male | 59 | IMF; BVVL; PUS | 2.750 | Male | 57 | Low value of RBC, HGB, HCT, MCV, MCH | 9.012 | Female | 53 | NA | 2.880 |
| Female | 79 | IMF; BVVL; PUS | 3.256 | Female | 77 | Low value of RBC, HGB, HCT, MCV, MCH | 3.696 |  |  |  |  |
| Male | 7 | IMF; BVVL; PUS | 3.179 | Male | 5 | Low value of RBC, HGB, HCT, MCV, MCH | 2.724 |  |  |  |  |
| Female | 81 | IMF; BVVL; PUS | 3.707 | Female | 79 | Low value of RBC, HGB, HCT, MCV, MCH | 3.360 |  |  |  |  |
| Male | 89 | IMF; BVVL; PUS | 2.992 | Male | 87 | Low value of RBC, HGB, HCT, MCV, MCH | 9.300 |  |  |  |  |
| Male | 68 | IMF; BVVL; PUS | 3.553 | Male | 66 | Low value of RBC, HGB, HCT, MCV, MCH | 2.268 |  |  |  |  |
| Male | 77 | IMF; BVVL; PUS | 4.213 | Female | 75 | Low value of RBC, HGB, HCT, MCV, MCH | 4.116 |  |  |  |  |
| Male | 75 | IMF; BVVL; PUS | 3.641 | Female | 73 | Low value of RBC, HGB, HCT, MCV, MCH | 2.592 |  |  |  |  |
| Male | 50 | IMF; BVVL; PUS | 2.013 | Male | 48 | Low value of RBC, HGB, HCT, MCV, MCH | 5.304 |  |  |  |  |
| Male | 74 | IMF; BVVL; PUS | 2.123 | Male | 72 | Low value of RBC, HGB, HCT, MCV, MCH | 2.640 |  |  |  |  |
| Male | 59 | IMF; BVVL; PUS | 2.101 | Male | 57 | Low value of RBC, HGB, HCT, MCV, MCH | 2.340 |  |  |  |  |
| Female | 65 | IMF; BVVL; PUS | 3.509 | Female | 63 | Low value of RBC, HGB, HCT, MCV, MCH | 4.920 |  |  |  |  |
| Female | 81 | IMF; BVVL; PUS | 2.684 | Female | 79 | Low value of RBC, HGB, HCT, MCV, MCH | 1.992 |  |  |  |  |
| Male | 76 | IMF; BVVL; PUS | 3.564 | Male | 74 | Low value of RBC, HGB, HCT, MCV, MCH | 9.864 |  |  |  |  |
| Female | 65 | IMF; BVVL; PUS | 2.134 | Male | 63 | Low value of RBC, HGB, HCT, MCV, MCH | 8.724 |  |  |  |  |
| Female | 28 | IMF; BVVL; PUS | 3.564 | Male | 26 | Low value of RBC, HGB, HCT, MCV, MCH | 3.792 |  |  |  |  |
| Male | 48 | IMF; BVVL; PUS | 3.201 | Male | 46 | Low value of RBC, HGB, HCT, MCV, MCH | 1.824 |  |  |  |  |
| Female | 56 | IMF; BVVL; PUS | 3.256 | Female | 54 | Low value of RBC, HGB, HCT, MCV, MCH | 2.028 |  |  |  |  |
| Male | 69 | IMF; BVVL; PUS | 3.773 | Male | 67 | Low value of RBC, HGB, HCT, MCV, MCH | 2.544 |  |  |  |  |
| Female | 61 | IMF; BVVL; PUS | 4.323 | Male | 59 | Low value of RBC, HGB, HCT, MCV, MCH | 3.972 |  |  |  |  |
| Female | 78 | IMF; BVVL; PUS | 4.202 | Male | 76 | Low value of RBC, HGB, HCT, MCV, MCH | 3.960 |  |  |  |  |
| Male | 48 | IMF; BVVL; PUS | 4.873 | Female | 46 | Low value of RBC, HGB, HCT, MCV, MCH | 6.396 |  |  |  |  |
| Male | 71 | IMF; BVVL; PUS | 4.169 | Female | 69 | Low value of RBC, HGB, HCT, MCV, MCH | 2.700 |  |  |  |  |
| Female | 84 | IMF; BVVL; PUS | 2.090 | Female | 82 | Low value of RBC, HGB, HCT, MCV, MCH | 2.628 |  |  |  |  |
| Male | 83 | IMF; BVVL; PUS | 3.465 | Female | 81 | Low value of RBC, HGB, HCT, MCV, MCH | 2.208 |  |  |  |  |
| Male | 68 | IMF; BVVL; PUS | 6.281 | Female | 66 | Low value of RBC, HGB, HCT, MCV, MCH | 2.748 |  |  |  |  |
| Female | 57 | IMF; BVVL; PUS | 4.257 | Female | 55 | Low value of RBC, HGB, HCT, MCV, MCH | 3.708 |  |  |  |  |
| Female | 60 | IMF; BVVL; PUS | 6.248 | Female | 58 | Low value of RBC, HGB, HCT, MCV, MCH | 3.048 |  |  |  |  |
| Female | 63 | IMF; BVVL; PUS | 3.652 | Male | 61 | Low value of RBC, HGB, HCT, MCV, MCH | 5.280 |  |  |  |  |
| Female | 88 | IMF; BVVL; PUS | 4.092 | Male | 86 | Low value of RBC, HGB, HCT, MCV, MCH | 6.396 |  |  |  |  |
| Male | 78 | IMF; BVVL; PUS | 5.093 | Male | 76 | Low value of RBC, HGB, HCT, MCV, MCH | 4.404 |  |  |  |  |
| Male | 80 | IMF; BVVL; PUS | 4.290 | Male | 78 | Low value of RBC, HGB, HCT, MCV, MCH | 6.624 |  |  |  |  |
| Male | 48 | IMF; BVVL; PUS | 3.509 | Male | 46 | Low value of RBC, HGB, HCT, MCV, MCH | 2.700 |  |  |  |  |
| Male | 64 | IMF; BVVL; PUS | 3.047 | Male | 62 | Low value of RBC, HGB, HCT, MCV, MCH | 2.568 |  |  |  |  |
| Female | 68 | IMF; BVVL; PUS | 3.938 | Female | 66 | Low value of RBC, HGB, HCT, MCV, MCH | 3.552 |  |  |  |  |
| Male | 73 | IMF; BVVL; PUS | 2.156 | Male | 71 | Low value of RBC, HGB, HCT, MCV, MCH | 2.364 |  |  |  |  |
| Male | 78 | IMF; BVVL; PUS | 4.818 | Female | 76 | Low value of RBC, HGB, HCT, MCV, MCH | 3.360 |  |  |  |  |
| Male | 22 | IMF; BVVL; PUS | 5.104 | Female | 20 | Low value of RBC, HGB, HCT, MCV, MCH | 3.852 |  |  |  |  |
| Female | 64 | IMF; BVVL; PUS | 4.796 | Female | 62 | Low value of RBC, HGB, HCT, MCV, MCH | 3.012 |  |  |  |  |
| Male | 69 | IMF; BVVL; PUS | 1.782 | Female | 67 | Low value of RBC, HGB, HCT, MCV, MCH | 2.844 |  |  |  |  |
| Male | 88 | IMF; BVVL; PUS | 2.277 | Female | 86 | Low value of RBC, HGB, HCT, MCV, MCH | 2.388 |  |  |  |  |
| Female | 48 | IMF; BVVL; PUS | 4.829 | Female | 46 | Low value of RBC, HGB, HCT, MCV, MCH | 5.136 |  |  |  |  |
| Male | 66 | IMF; BVVL; PUS | 4.136 | Male | 64 | Low value of RBC, HGB, HCT, MCV, MCH | 6.828 |  |  |  |  |
| Female | 72 | IMF; BVVL; PUS | 3.520 | Female | 70 | Low value of RBC, HGB, HCT, MCV, MCH | 5.136 |  |  |  |  |
| Female | 89 | IMF; BVVL; PUS | 3.784 | Female | 87 | Low value of RBC, HGB, HCT, MCV, MCH | 2.196 |  |  |  |  |
| Female | 60 | IMF; BVVL; PUS | 3.861 | Female | 58 | Low value of RBC, HGB, HCT, MCV, MCH | 5.292 |  |  |  |  |
| Male | 73 | IMF; BVVL; PUS | 3.366 | Male | 71 | Low value of RBC, HGB, HCT, MCV, MCH | 5.484 |  |  |  |  |
| Female | 89 | IMF; BVVL; PUS | 2.750 | Female | 87 | Low value of RBC, HGB, HCT, MCV, MCH | 2.760 |  |  |  |  |
| Male | 62 | IMF; BVVL; PUS | 4.279 | Male | 60 | Low value of RBC, HGB, HCT, MCV, MCH | 5.820 |  |  |  |  |
| Female | 88 | IMF; BVVL; PUS | 3.916 | Female | 86 | Low value of RBC, HGB, HCT, MCV, MCH | 2.916 |  |  |  |  |
| Male | 53 | IMF; BVVL; PUS | 3.674 | Female | 51 | Low value of RBC, HGB, HCT, MCV, MCH | 3.504 |  |  |  |  |
| Male | 29 | IMF; BVVL; PUS | 2.530 | Female | 27 | Low value of RBC, HGB, HCT, MCV, MCH | 2.088 |  |  |  |  |
| Female | 46 | IMF; BVVL; PUS | 3.531 | Female | 44 | Low value of RBC, HGB, HCT, MCV, MCH | 2.028 |  |  |  |  |
| Female | 76 | IMF; BVVL; PUS | 5.027 | Female | 74 | Low value of RBC, HGB, HCT, MCV, MCH | 8.808 |  |  |  |  |
| Male | 79 | IMF; BVVL; PUS | 3.652 | Male | 77 | Low value of RBC, HGB, HCT, MCV, MCH | 3.780 |  |  |  |  |
| Male | 47 | IMF; BVVL; PUS | 4.015 | Male | 45 | Low value of RBC, HGB, HCT, MCV, MCH | 3.624 |  |  |  |  |
| Female | 77 | IMF; BVVL; PUS | 3.146 | Male | 75 | Low value of RBC, HGB, HCT, MCV, MCH | 2.760 |  |  |  |  |
| Female | 54 | IMF; BVVL; PUS | 5.093 | Female | 52 | Low value of RBC, HGB, HCT, MCV, MCH | 5.988 |  |  |  |  |
| Male | 44 | IMF; BVVL; PUS | 3.355 | Female | 42 | Low value of RBC, HGB, HCT, MCV, MCH | 5.448 |  |  |  |  |
| Male | 62 | IMF; BVVL; PUS | 4.488 | Male | 60 | Low value of RBC, HGB, HCT, MCV, MCH | 5.436 |  |  |  |  |
| Male | 82 | IMF; BVVL; PUS | 3.861 | Male | 80 | Low value of RBC, HGB, HCT, MCV, MCH | 4.872 |  |  |  |  |
| Female | 85 | IMF; BVVL; PUS | 4.125 | Female | 83 | Low value of RBC, HGB, HCT, MCV, MCH | 3.156 |  |  |  |  |
| Male | 82 | IMF; BVVL; PUS | 3.839 | Male | 80 | Low value of RBC, HGB, HCT, MCV, MCH | 4.428 |  |  |  |  |
| Male | 84 | IMF; BVVL; PUS | 3.938 | Female | 82 | Low value of RBC, HGB, HCT, MCV, MCH | 3.780 |  |  |  |  |
| Male | 84 | IMF; BVVL; PUS | 2.508 | Female | 82 | Low value of RBC, HGB, HCT, MCV, MCH | 3.036 |  |  |  |  |
| Male | 77 | IMF; BVVL; PUS | 4.356 | Male | 75 | Low value of RBC, HGB, HCT, MCV, MCH | 3.972 |  |  |  |  |
| Female | 57 | IMF; BVVL; PUS | 3.850 | Female | 55 | Low value of RBC, HGB, HCT, MCV, MCH | 3.540 |  |  |  |  |
| Male | 83 | IMF; BVVL; PUS | 9.801 | Male | 81 | Low value of RBC, HGB, HCT, MCV, MCH | 1.824 |  |  |  |  |
| Female | 65 | IMF; BVVL; PUS | 3.410 | Female | 63 | Low value of RBC, HGB, HCT, MCV, MCH | 1.716 |  |  |  |  |
| Male | 83 | IMF; BVVL; PUS | 2.871 | Female | 81 | Low value of RBC, HGB, HCT, MCV, MCH | 9.864 |  |  |  |  |
| Male | 79 | IMF; BVVL; PUS | 3.421 | Female | 77 | Low value of RBC, HGB, HCT, MCV, MCH | 3.408 |  |  |  |  |
| Male | 66 | IMF; BVVL; PUS | 5.302 | Male | 64 | Low value of RBC, HGB, HCT, MCV, MCH | 4.656 |  |  |  |  |
| Male | 27 | IMF; BVVL; PUS | 2.904 | Female | 25 | Low value of RBC, HGB, HCT, MCV, MCH | 3.492 |  |  |  |  |
| Male | 61 | IMF; BVVL; PUS | 4.532 | Male | 59 | Low value of RBC, HGB, HCT, MCV, MCH | 4.968 |  |  |  |  |
| Male | 73 | IMF; BVVL; PUS | 5.412 | Male | 71 | Low value of RBC, HGB, HCT, MCV, MCH | 7.656 |  |  |  |  |
| Male | 62 | IMF; BVVL; PUS | 2.871 | Male | 60 | Low value of RBC, HGB, HCT, MCV, MCH | 11.304 |  |  |  |  |
| Male | 62 | IMF; BVVL; PUS | 3.740 | Male | 60 | Low value of RBC, HGB, HCT, MCV, MCH | 3.420 |  |  |  |  |
| Male | 31 | IMF; BVVL; PUS | 2.255 | Male | 29 | Low value of RBC, HGB, HCT, MCV, MCH | 6.936 |  |  |  |  |
| Male | 62 | IMF; BVVL; PUS | 2.651 | Female | 60 | Low value of RBC, HGB, HCT, MCV, MCH | 4.968 |  |  |  |  |
| Male | 77 | IMF; BVVL; PUS | 2.475 | Female | 75 | Low value of RBC, HGB, HCT, MCV, MCH | 6.756 |  |  |  |  |
| Male | 41 | IMF; BVVL; PUS | 2.464 | Male | 39 | Low value of RBC, HGB, HCT, MCV, MCH | 8.784 |  |  |  |  |
| Male | 86 | IMF; BVVL; PUS | 2.794 | Male | 84 | Low value of RBC, HGB, HCT, MCV, MCH | 7.656 |  |  |  |  |
| Male | 67 | IMF; BVVL; PUS | 2.134 | Male | 65 | Low value of RBC, HGB, HCT, MCV, MCH | 3.372 |  |  |  |  |
| Male | 69 | IMF; BVVL; PUS | 2.332 | Male | 70 | Low value of RBC, HGB, HCT, MCV, MCH | 2.400 |  |  |  |  |
| Male | 62 | IMF; BVVL; PUS | 3.025 | Male | 25 | Low value of RBC, HGB, HCT, MCV, MCH | 7.068 |  |  |  |  |
| Male | 82 | IMF; BVVL; PUS | 2.332 | Female | 58 | Low value of RBC, HGB, HCT, MCV, MCH | 6.756 |  |  |  |  |
| Female | 83 | IMF; BVVL; PUS | 4.785 | Female | 48 | Low value of RBC, HGB, HCT, MCV, MCH | 10.884 |  |  |  |  |
| Female | 45 | IMF; BVVL; PUS | 5.830 | Male | 78 | Low value of RBC, HGB, HCT, MCV, MCH | 7.320 |  |  |  |  |
| Female | 42 | IMF; BVVL; PUS | 4.730 | Male | 54 | Low value of RBC, HGB, HCT, MCV, MCH | 6.936 |  |  |  |  |
| Male | 64 | IMF; BVVL; PUS | 3.850 | Male | 65 | Low value of RBC, HGB, HCT, MCV, MCH | 3.120 |  |  |  |  |
| Female | 27 | IMF; BVVL; PUS | 6.710 | Female | 54 | Low value of RBC, HGB, HCT, MCV, MCH | 2.184 |  |  |  |  |
| Female | 38 | IMF; BVVL; PUS | 6.160 | Female | 19 | Low value of RBC, HGB, HCT, MCV, MCH | 4.044 |  |  |  |  |
| Female | 79 | IMF; BVVL; PUS | 4.785 | Female | 18 | Low value of RBC, HGB, HCT, MCV, MCH | 2.184 |  |  |  |  |
| Female | 62 | IMF; BVVL; PUS | 3.916 | Male | 40 | Low value of RBC, HGB, HCT, MCV, MCH | 4.476 |  |  |  |  |
| Female | 45 | IMF; BVVL; PUS | 3.597 | Male | 66 | Low value of RBC, HGB, HCT, MCV, MCH | 2.880 |  |  |  |  |
| Male | 84 | IMF; BVVL; PUS | 3.773 | Female | 37 | Low value of RBC, HGB, HCT, MCV, MCH | 3.060 |  |  |  |  |
| Male | 32 | IMF; BVVL; PUS | 2.827 | Female | 28 | Low value of RBC, HGB, HCT, MCV, MCH | 2.760 |  |  |  |  |
| Female | 16 | IMF; BVVL; PUS | 2.376 | Female | 54 | Low value of RBC, HGB, HCT, MCV, MCH | 4.044 |  |  |  |  |
| Male | 24 | IMF; BVVL; PUS | 2.651 | Female | 67 | Low value of RBC, HGB, HCT, MCV, MCH | 2.184 |  |  |  |  |
| Male | 21 | IMF; BVVL; PUS | 3.069 | Female | 63 | Low value of RBC, HGB, HCT, MCV, MCH | 4.476 |  |  |  |  |
| Female | 20 | IMF; BVVL; PUS | 2.541 | Female | 83 | Low value of RBC, HGB, HCT, MCV, MCH | 2.880 |  |  |  |  |
| Female | 42 | IMF; BVVL; PUS | 3.058 | Female | 84 | Low value of RBC, HGB, HCT, MCV, MCH | 3.060 |  |  |  |  |
| Male | 68 | IMF; BVVL; PUS | 4.906 | Female | 46 | Low value of RBC, HGB, HCT, MCV, MCH | 2.760 |  |  |  |  |
| Male | 39 | IMF; BVVL; PUS | 5.456 | Female | 43 | Low value of RBC, HGB, HCT, MCV, MCH | 2.124 |  |  |  |  |
| Female | 30 | IMF; BVVL; PUS | 7.062 | Female | 65 | Low value of RBC, HGB, HCT, MCV, MCH | 2.256 |  |  |  |  |
| Female | 56 | IMF; BVVL; PUS | 6.270 | Male | 28 | Low value of RBC, HGB, HCT, MCV, MCH | 2.640 |  |  |  |  |
| Female | 72 | IMF; BVVL; PUS | 6.358 | Male | 39 | Low value of RBC, HGB, HCT, MCV, MCH | 2.952 |  |  |  |  |
| Male | 27 | IMF; BVVL; PUS | 7.348 | Female | 80 | Low value of RBC, HGB, HCT, MCV, MCH | 2.268 |  |  |  |  |
| Male | 60 | IMF; BVVL; PUS | 5.170 | Female | 63 | Low value of RBC, HGB, HCT, MCV, MCH | 3.432 |  |  |  |  |
| Female | 50 | IMF; BVVL; PUS | 5.456 | Female | 46 | Low value of RBC, HGB, HCT, MCV, MCH | 4.092 |  |  |  |  |
| Female | 80 | IMF; BVVL; PUS | 4.653 | Female | 85 | Low value of RBC, HGB, HCT, MCV, MCH | 2.460 |  |  |  |  |
| Male | 56 | IMF; BVVL; PUS | 4.587 | Male | 33 | Low value of RBC, HGB, HCT, MCV, MCH | 1.860 |  |  |  |  |
| Male | 67 | IMF; PUS | 5.643 | Female | 17 | Low value of RBC, HGB, HCT, MCV, MCH | 9.636 |  |  |  |  |
|  |  |  |  | Female | 25 | Low value of RBC, HGB, HCT, MCV, MCH | 2.784 |  |  |  |  |

IS: Ischemic stroke; IDA: Iron-deficiency anemia; Tf: Transferrin; NA: Not applicable;

RBC: Red blood cell; HGB: Hemoglobin; HCT: Hematocrit; MCV: Mean corpusular volume; MCH: Mean corpusular hemoglobin;

IMF: Inability to move or feel; PUS: Problems of understanding and speaking; BVVL: Blurring of vision or vision loss.

**Table S2** Clinical features, and CSF Tf concentrations in patients with IS, and healthy controls.

| IS | | | | Controls (Normal physical examinees) | | | |
| --- | --- | --- | --- | --- | --- | --- | --- |
| Gender | Age (years) | Main clinical features | Tf (mg/L) | Gender | Age (years) | Main clinical features | Tf (mg/L) |
| Female | 81 | IMF; BVVL | 12.900 | Female | 49 | NA | 9.900 |
| Male | 87 | IMF; BVVL; PUS | 10.900 | Female | 70 | NA | 15.600 |
| Female | 80 | BVVL; PUS | 15.700 | Female | 80 | NA | 6.800 |
| Male | 75 | IMF; BVVL; PUS | 15.600 | Male | 45 | NA | 17.900 |
| Female | 67 | IMF; BVVL; PUS | 13.700 | Female | 61 | NA | 15.200 |
| Female | 73 | IMF; BVVL; PUS | 17.400 | Male | 67 | NA | 10.500 |
| Male | 69 | IMF; BVVL | 20.500 | Male | 79 | NA | 11.500 |
| Female | 71 | IMF; PUS | 17.200 | Male | 45 | NA | 7.900 |
| Male | 51 | IMF; BVVL; PUS | 28.900 | Male | 37 | NA | 16.900 |
| Male | 66 | IMF; PUS | 15.400 | Male | 75 | NA | 12.300 |
| Male | 63 | IMF; PUS | 25.700 | Female | 39 | NA | 13.400 |
| Male | 51 | IMF; BVVL; PUS | 17.500 | Male | 48 | NA | 15.700 |
| Male | 65 | IMF; BVVL; PUS | 26.700 | Female | 70 | NA | 20.600 |
| Male | 48 | IMF; BVVL; PUS | 19.900 | Female | 51 | NA | 12.400 |
| Male | 51 | IMF; BVVL | 30.200 | Male | 38 | NA | 10.900 |
| Male | 66 | IMF; BVVL; PUS | 20.500 | Female | 30 | NA | 19.900 |
| Female | 76 | IMF; BVVL; PUS | 20.400 | Male | 42 | NA | 12.400 |
| Male | 63 | IMF; BVVL | 26.900 | Male | 29 | NA | 12.900 |
| Female | 66 | IMF; BVVL; PUS | 19.700 | Female | 80 | NA | 14.900 |
| Female | 54 | IMF; PUS | 28.000 | Female | 47 | NA | 15.600 |
| Female | 68 | IMF; BVVL; PUS | 15.600 | Male | 77 | NA | 12.300 |
| Male | 51 | IMF; BVVL; PUS | 12.900 | Male | 41 | NA | 13.800 |
| Female | 54 | IMF; BVVL; PUS | 12.700 | Male | 44 | NA | 9.900 |
| Female | 48 | IMF; BVVL; PUS | 16.500 | Male | 31 | NA | 15.300 |
| Female | 79 | IMF; BVVL; PUS | 12.900 | Female | 82 | NA | 14.100 |
| Female | 64 | IMF; BVVL; PUS | 17.400 | Male | 49 | NA | 10.500 |
| Male | 69 | IMF; BVVL; PUS | 20.500 | Male | 82 | NA | 13.100 |
| Female | 47 | IMF; BVVL; PUS | 16.200 | Male | 47 | NA | 12.900 |
| Male | 84 | IMF; BVVL; PUS | 27.500 | Female | 63 | NA | 13.500 |
| Male | 90 | IMF; BVVL; PUS | 15.400 | Female | 69 | NA | 12.300 |
| Male | 66 | IMF; BVVL; PUS | 25.700 | Female | 81 | NA | 12.600 |
| Female | 51 | IMF; BVVL; PUS | 17.500 | Male | 47 | NA | 14.700 |
| Female | 83 | IMF; BVVL; PUS | 21.900 | Female | 39 | NA | 18.900 |
| Female | 78 | IMF; BVVL; PUS | 18.800 | Female | 53 | NA | 11.400 |
| Male | 70 | IMF; BVVL; PUS | 31.700 | Female | 69 | NA | 13.100 |
| Male | 76 | IMF; BVVL; PUS | 20.500 | Female | 81 | NA | 12.900 |
| Female | 72 | IMF; BVVL; PUS | 22.500 | Male | 47 | NA | 19.400 |
| Male | 74 | IMF; BVVL; PUS | 28.900 | Female | 39 | NA | 14.900 |
| Male | 54 | IMF; BVVL; PUS | 16.700 | Female | 53 | NA | 12.000 |
| Male | 69 | IMF; BVVL; PUS | 29.000 | Male | 72 | NA | 15.900 |

IS: Ischemic stroke; CSF: Cerebrospinal fluid; Tf: Transferrin; NA: Not applicable;

IMF: Inability to move or feel; PUS: Problems of understanding and speaking; BVVL: Blurring of vision or vision loss.

**Table S3. Clinical features and plasma Tf concentrations in women VTE patients using CC**

| Gender | Age (years) | Main clinical features | Tf (mg/ml) |
| --- | --- | --- | --- |
| VTE patients | | | |
| Female | 52 | VVTE | 6.440 |
| Female | 43 | VVTE | 4.923 |
| Female | 37 | VVTE | 3.867 |
| Female | 39 | VVTE | 3.814 |
| Female | 35 | VVTE | 6.230 |
| Female | 54 | VVTE | 4.369 |
| Female | 55 | VVTE | 3.828 |
| Female | 40 | VVTE | 4.540 |
| Female | 32 | VVTE | 5.293 |
| Female | 56 | VVTE | 6.798 |
| Female | 49 | VVTE | 5.016 |
| Female | 41 | VVTE | 4.910 |
| Female | 52 | VVTE | 5.623 |
| Female | 38 | VVTE | 3.841 |
| Female | 35 | VVTE | 2.640 |
| Female | 45 | VVTE | 4.950 |
| Female | 36 | VVTE | 5.385 |
| Female | 34 | VVTE | 5.940 |
| Female | 39 | VVTE | 4.224 |
| Female | 50 | VVTE | 10.084 |
| Female | 39 | VVTE | 7.194 |
| Female | 31 | VVTE | 3.498 |
| Female | 47 | VVTE | 4.844 |
| Female | 35 | VVTE | 4.752 |
| Female | 39 | VVTE | 7.893 |
| Female | 40 | VVTE | 6.349 |
| Female | 44 | VVTE | 4.171 |
| Female | 43 | VVTE | 6.336 |
| Female | 55 | VVTE | 5.900 |
| Female | 38 | VVTE | 4.527 |
| Healthy people | | | |
| Female | 45 | NA | 2.976 |
| Female | 33 | NA | 3.408 |
| Female | 43 | NA | 2.268 |
| Female | 49 | NA | 1.692 |
| Female | 39 | NA | 4.620 |
| Female | 42 | NA | 3.984 |
| Female | 37 | NA | 2.676 |
| Female | 45 | NA | 2.952 |
| Female | 42 | NA | 2.052 |
| Female | 45 | NA | 2.472 |
| Female | 39 | NA | 2.436 |
| Female | 48 | NA | 2.160 |
| Female | 34 | NA | 4.320 |
| Female | 54 | NA | 2.268 |
| Female | 38 | NA | 4.344 |
| Female | 29 | NA | 2.904 |
| Female | 46 | NA | 4.464 |
| Female | 37 | NA | 2.160 |
| Female | 27 | NA | 2.700 |
| Female | 45 | NA | 3.828 |
| Female | 45 | NA | 2.880 |
| Female | 32 | NA | 3.120 |
| Female | 51 | NA | 2.928 |
| Female | 47 | NA | 3.876 |
| Female | 51 | NA | 5.160 |
| Female | 45 | NA | 3.000 |
| Female | 36 | NA | 3.360 |
| Female | 39 | NA | 2.280 |
| Female | 47 | NA | 3.912 |
| Female | 29 | NA | 2.388 |

VTE: venous thromboembolism; CC: combined oral contraceptives; Tf: transferrin; VVTE: visible venous thromboembolism; NA: Not Applicable;

**Table S4** Blood routine examination.

|  | RBC (10^12^/L) | HGB (g/L) | HCT (%) | MCV (fL) | MCH (pg) |
| --- | --- | --- | --- | --- | --- |
| Normal mice | 8.49 (1.310) | 122 (3.456) | 39.7 (4.561) | 46.8 (2.134) | 14.3 (1.230) |
| Low-iron induced mice | 7.89 (2.340) | 84 (5.655) | 26.4 (3.231) | 33.5 (3.913) | 10.6 (0.985) |
| High-iron induced mice | 8.29 (1.572) | 111 (2.345) | 35.9 (2.345) | 46.8 (4.560) | 14.4 (1.113) |

Routine blood tests (including RBC, HGB, HCT, MCV, MCH) of mice induced by normal forage, low-iron forage (Fe < 10 mg/kg), high-iron forage (Fe > 90 mg/kg) were examined. Data represent mean (SD), n=10. RBC: Red blood cell; HGB: Hemoglobin; HCT: Hematocrit; MCV: Mean corpusular volume; MCH: Mean corpusular hemoglobin.
